# Angiotensin Converting Enzyme (ACE) expression in microglia reduces amyloid β deposition and neurodegeneration by increasing SYK signaling and endolysosomal trafficking

**DOI:** 10.1101/2024.04.24.590837

**Authors:** Andrew R. Gomez, Hyae Ran Byun, Shaogen Wu, AKM Ghulam Muhammad, Jasmine Ikbariyeh, Jaelin Chen, Alek Muro, Lin Li, Kenneth E. Bernstein, Richard Ainsworth, Warren G. Tourtellotte

**Author notes:** Correspondence to: Warren G. Tourtellotte, MD, PhD, FCAP Department of Pathology and Laboratory Medicine Cedars-Sinai Medical Center 8700 Beverly Blvd., Rm. 8719 Los Angeles, CA 90048.

## Abstract

Genome-wide association studies (GWAS) have identified many gene polymorphisms associated with an increased risk of developing Late Onset Alzheimer’s Disease (LOAD). Many of these LOAD risk-associated alleles alter disease pathogenesis by influencing microglia innate immune responses and lipid metabolism. Angiotensin Converting Enzyme (ACE), a GWAS LOAD risk-associated gene best known for its role in regulating systemic blood pressure, also enhances innate immunity and lipid processing in peripheral myeloid cells, but a role for ACE in modulating the function of myeloid-derived microglia remains unexplored. Using novel mice engineered to express ACE in microglia and CNS associated macrophages (CAMs), we find that ACE expression in microglia reduces Aβ plaque load, preserves vulnerable neurons and excitatory synapses, and greatly reduces learning and memory abnormalities in the 5xFAD amyloid mouse model of Alzheimer’s Disease (AD). ACE-expressing microglia show enhanced Aβ phagocytosis and endolysosomal trafficking, increased clustering around amyloid plaques, and increased SYK tyrosine kinase activation downstream of the major Aβ receptors, TREM2 and CLEC7A. Single microglia sequencing and digital spatial profiling identifies downstream SYK signaling modules that are expressed by ACE expression in microglia that mediate endolysosomal biogenesis and trafficking, mTOR and PI3K/AKT signaling, and increased oxidative phosphorylation, while gene silencing or pharmacologic inhibition of SYK activity in ACE-expressing microglia abrogates the potentiated Aβ engulfment and endolysosomal trafficking. These findings establish a role for ACE in enhancing microglial immune function and they identify a potential use for ACE-expressing microglia as a cell-based therapy to augment endogenous microglial responses to Aβ in AD.

## INTRODUCTION

Alzheimer’s Disease (AD) is the leading cause of dementia and a progressive neurodegenerative disease which currently has no cure (1). AD is characterized by intraneuronal aggregates of paired helical filaments formed from hyper-phosphorylated microtubule associated protein tau (neurofibrillary tangles) and abnormally cleaved Amyloid Precursor Protein (APP) that aggregates into soluble amyloid-beta (Aβ) oligomers and insoluble macromers that form extracellular plaques in the brain (2). Together, these neuropathological changes lead to neurodegeneration and cognitive decline which is distinct from normal aging. Microglia, the myeloid-derived resident innate immune phagocytes of the central nervous system, have an important role in clearing protein aggregates and maintaining amyloid proteostasis in the brain (3, 4). While germline mutations in genes such as Presenilin and APP account for a relatively small number of early onset cases of familial AD (5), Genome-Wide Association Studies (GWAS) have provided accumulating evidence that sequence polymorphisms can influence gene expression and coding sequences to confer increased risk of the more common Late-Onset AD (LOAD) (6–8). Interestingly, many of these genes encode proteins that appear to modulate the ability of microglia to recognize, engulf, traffic and degrade protein aggregates such as Aβ (9). While a detailed understanding of the convergent signaling pathways within microglia that are altered by LOAD polymorphisms is only beginning to emerge, they hold promise for discovering effective pharmacologic or cell-based therapies for AD.

Angiotensin Converting Enzyme (ACE) has been repeatedly implicated as a LOAD gene, but how it functions in AD pathogenesis is poorly understood (6, 7, 10–14). ACE is a dipeptidyl carboxypeptidase that is best known for its role in blood pressure regulation where it cleaves angiotensin I (AngI) into angiotensin II (AngII), a potent vasoconstrictor (15). Evidence suggests that ACE catalytic function may have a protective effect on the development of dementia considering that ACE function modifying anti-hypertensive drugs appear to decrease the overall rate of dementia (16–18), but more specifically ACE inhibitor drugs that reduce blood pressure by blocking the AngII receptor, AGTR1 rather than its catalytic activity, appear to be most effective at decreasing the rate of dementia (19–21). ACE is a relatively promiscuous peptidase which cleaves substrates not involved in blood pressure regulation (22), suggesting that its protective role in AD may be due to non-blood pressure regulating functions such as in myeloid cell development, myeloid immune function and cleavage of neurotoxic Aβ_1-42_ protein to less neurotoxic Aβ_1-40_ protein in AD (23–25). In mice, a clinical dose of ACE inhibitor that blocks its catalytic activity worsened amyloid pathology in the hAPPSwInd amyloid AD model, as did induced ACE haploinsufficiency (26). Interestingly, ACE expression in myeloid-derived macrophages and granulocytes leads to a profound enhancement of their bactericidal, anti-tumor and anti-atherogenic activity (27–29). Thus, ACE may have a comparable role in potentiating myeloid-derived microglia function like it does in peripheral myeloid-derived macrophages and neutrophils.

Here, we tested the hypothesis that ACE expression in microglia can modulate their ability to process Aβ and to alter the course of neurodegeneration and cognitive decline in an amyloid model of AD. We generated a novel mouse model which expresses ACE in microglia and CNS-associated macrophages (CAMs) to interrogate whether it can alter microglia function and the molecular, histopathologic and cognitive hallmarks of AD that have previously been well-characterized in the 5xFAD amyloid mouse model of AD. We found that ACE-expressing microglia have enhanced ability to migrate toward plaques and engulf neurotoxic Aβ_42_, and to reduce vulnerable excitatory synapse loss, neurodegeneration and cognitive abnormalities associated with Aβ deposition in the brain over time. Interestingly, we found in vivo and in vitro evidence that ACE expression in microglia increases SYK tyrosine phosphorylation and activity which is known to mediate signaling downstream of the major microglia Aβ receptors, TREM2 and CLEC7A, and to orchestrate the microglia immune response to Aβ (30–32). Taken together, these results identify a role for ACE that enhances Aβ processing in microglia to protect the brain from neurodegeneration.

## RESULTS

### Expression of ACE in microglia

Endogenous ACE expression is not detectable in homeostatic microglia or in microglia activated around amyloid plaques. However, in ACE^10/10^ transgenic mice that express ACE in peripheral macrophages (27), there is also strong expression of ACE in microglia (Fig. S1), suggesting that the previous results showing decreased amyloid burden and behavioral abnormalities in ACE^10/10^::APP/PS1 mice may have primarily been a result of ACE function in microglia rather than macrophages (33). To examine the function of ACE specifically in microglia, novel mice were generated by targeting the safe harbor Rosa locus with a construct regulated by the CAG promoter disrupted by a loxP flanked transcription stop (LSL) sequence upstream of a self-cleaving 2A peptide for TurboGFP and hACE-3xF proteins (34, 35) (Fig. S2A). These novel Rosa^+/hACE-flox^ mice were mated to Cx3Cr1^+/CreERT2^ mice that express Tamoxifen (TMX)-dependent cre-recombinase in myeloid cells (36) to generate RACE-(Rosa^+/+^;Cx3Cr1^+/CreERT2^) and RACE+ (Rosa^+/hACE-flox^;Cx3Cr1^+/CreERT2^) mice (Fig. S2B). To restrict hACE expression to yolk sac-derived myeloid cells that engraft into the CNS during development (e.g., microglia and <u>C</u>NS-<u>A</u>ssociated <u>M</u>acrophages – CAMs, that consist of a small population of perivascular macrophages, dural macrophages, and choroid plexus macrophages) (37), RACE- and RACE+ mice received TMX injections (125 mg/kg, IP) on four consecutive days between postnatal days 28 to 31 to activate transgene expression in myeloid cells in RACE+ (hereafter referred to as <u>R+</u>), but not in RACE-(hereafter referred to as <u>R-</u>) mice (Fig. S2C). After TMX treatment, peripheral myeloid cells with loxP site recombination (transgene-ON) are replaced by turnover of non-loxP recombined (transgene-OFF) cells from the bone marrow approximately 21 days after TMX treatment, whereas myeloid-derived cells engrafted in the brain early during embryogenesis, such as microglia, do not turn over in the brain and retain their transgene expression (36, 38). As expected, 3 weeks after TMX injection, flow cytometry analysis of myeloid cells isolated from brain, blood and spleen showed that the TurboGFP transgene was highly expressed in brain myeloid cells from R+ mice, whereas transgene expression in peripheral blood and spleen myeloid cells returned to undetectable levels (Fig. S2D, E). hACE expression in R+ mice was restricted to microglia and CAMs (Fig. 1A), and it was localized within the cell body, cell surface and on delicate microglia filopodia not visualized by Iba1 staining (Fig 1B). hACE protein expression was restricted to >96% of microglia in R+ mice and it was undetectable in microglia from R-mice (Fig. S2F).

**Fig. 1.**
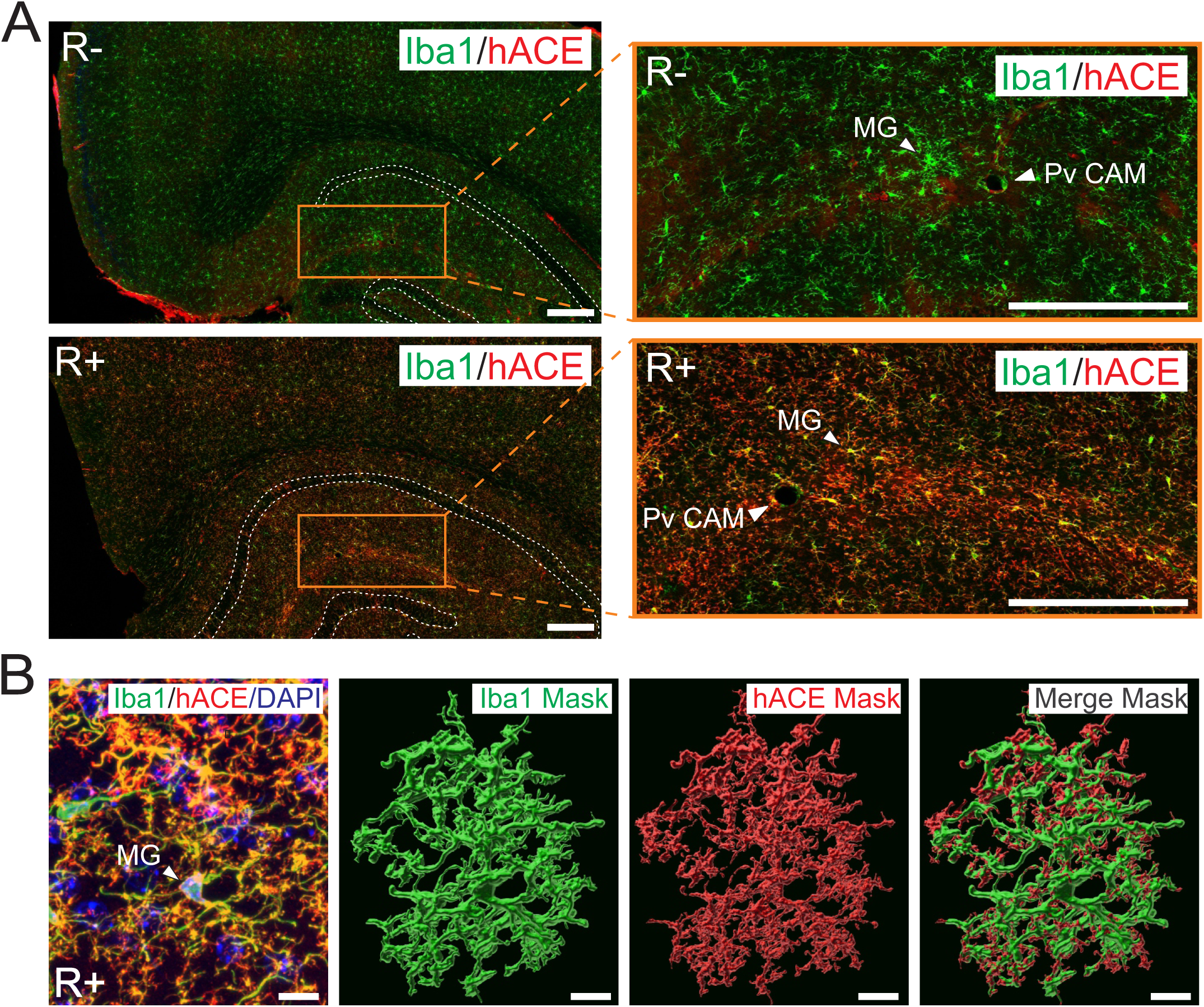
RACE+ mice treated with Tamoxifen (R+) express ACE in microglia. **(A)** Endogenous ACE expression was localized to vascular endothelial cells, but not microglia or Pv CAMs in R- mice. In R+ mice, ACE expression is upregulated in MG and CAMs throughout the brain. Microglia (MG) and perivascular CAM (Pv Cam) shown in Stratum Oriens of the hippocampus. **(B)** 3-dimensional reconstruction of microglia 50 µM thick tissue sections from R+ mice showed marked hACE expression restricted to the cell body, processes and filopodia not labeled by Iba1. (A, scale bar = 250 µM and B = 10 µM)

### ACE expression in microglia decreases Aβ_1-42_ amyloid plaque burden in 5xFAD mice

RACE and 5xFAD mice were mated to examine whether ACE expression in microglia can modulate their immune function and enhance their ability to reduce amyloid plaque burden in an amyloid model of AD. 5xFAD transgenic mice express 5 human mutations of APP and PS1, they accumulate Aβ_42_ protein that aggregates into extracellular amyloid plaques in the brain (39), and have neurodegeneration and cognitive abnormalities similar to AD (39, 40). 5xFAD hemizygous mice from the original Tg6799 transgenic line were backcrossed to C57BL6J (5xFAD^+/Tg6799^, hereafter referred to as AD+) (39), mated to RACE mice and injected with TMX to generate R-;AD- (Cx3Cr1^+/CreERT2^;Rosa^+/+^;5xFAD^+/+^), R+;AD- (Cx3Cr1^+/CreERT2^;Rosa^+/ACE-flox^;5xFAD^+/+^), R-;AD+ (Cx3Cr1^+/CreERT2^;Rosa^+/+^;5xFAD^+/Tg6799^), and R+;AD+ (Cx3Cr1^+/CreERT2^;Rosa^+/ACE-flox^;5xFAD^+/Tg6799^) mice. By 9 months of age, R-;AD+ mice consistently accumulated a large number of Aβ_42_ containing plaques throughout the brain (Fig. 2 A and A’) and in R+;AD+ mice, plaque density was markedly decreased throughout the brain (Fig. 2B and B’) and where quantified, it was decreased by 3.5-fold in the hippocampal CA1 region (p < 0.0001), 2.3-fold in somatosensory cortex (p < 0.001) and 2.6-fold in the amygdala (p< 0.001) (Fig 2C).

**Fig. 2.**
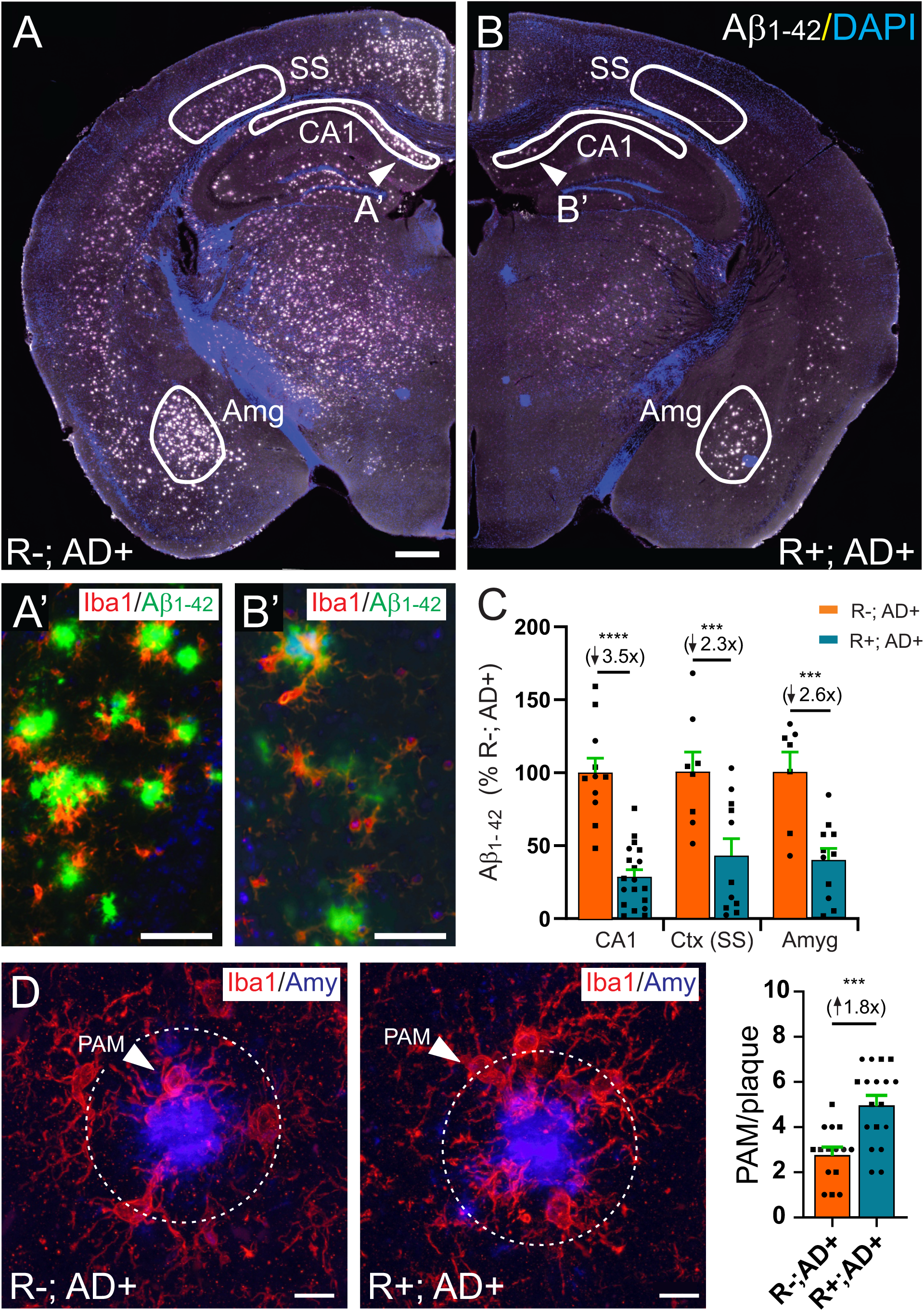
ACE expression in microglia reduced Aβ plaque burden in brains from 5xFAD mice. In 6-month-old mice, Aβ_1-42_ containing plaques were markedly decreased in **(B)** R+;AD+ compared to **(A)** R-;AD+ brains. **(A’, B’)** Aβ_1-42_ containing plaques (green) were surrounded by Iba1 expressing microglia (red). **(C)** Plaque density was reduced in CA1 of the hippocampus by 3.5-fold (p < 0.0001), in somatosensory cortex by 2.3-fold (p < 0.001) and in the amygdala by 2.6-fold (p < 0.001) in R+; AD+ compared to R-; AD+ brains. **(D)** Plaques were identified by staining them with Amylo-Glo (blue) and plaque associated microglia (PAM) labeled with an antibody to Iba1 were defined as Iba1+ microglia within 10 µM of a plaque edge. There were 1.8-fold more PAM surrounding plaques in R+;AD+ compared to R-;AD+ mice (p < 0.001). (A, B, scale bar = 500 µM, A’, B’ inset = 50 µM, D = 10 µM).

Injury, pathogens and other insults to the brain lead to microglia activation that is characterized in part by changes in their morphology and increased cytokine production (41). Under homeostatic conditions, microglia have highly branched and elongated processes that shorten and become less branched upon neuroinflammatory activation (42, 43). Appropriately reflecting their activated state, microglia had significantly decreased branching and process length in R-;AD+ mice compared to R-;AD-mice (p < 0.0001). Interestingly, ACE expression in microglia significantly increased process branching and length in R+;AD- mice (p < 0.0001) and it restored process branching and length to control (R-;AD-) levels in R+;AD+ mice (p < 0.05; Fig. S3A, B). Among nine pro-inflammatory cytokines examined, TNF-α (p < 0.0001), IL1-β (p<0.0001) and CXCL1 (p < 0.05) were elevated in R-;AD+ compared to R-;AD- mice. ACE expression in microglia did not significantly alter the expression of TNF-α or CXCL1, but it modestly increased expression of IL1-β in R+;AD+ compared to R-;AD+ mice (p < 0.01; Fig. S4). In addition, IL-12p70 and IL-5 were significantly decreased in R+;AD- mice compared to R-;AD- mice (p < 0.05), but unchanged in R-;AD+ or R+;AD+ mice compared to R-;AD- mice. ACE expression in microglia slightly decreased the ratio of neurotoxic Aβ_1-42_ to non-neurotoxic Aβ_1-40_ protein production in the brain by 1.2-fold (p< 0.05; Fig. S5). Thus, ACE expression in microglia significantly reduces plaque load in the brains of 5xFAD mice, and while it alters microglia morphology and slightly decreases neurotoxic Aβ_1-42_ production, it has very little effect on the production of the proinflammatory cytokines that were examined.

### ACE expression in microglia enhances their association with plaques and increases Aβ phagocytosis and endolysosomal trafficking

ACE expression in microglia increased their accumulation around plaques, leading to 1.8-fold more plaque-associated microglia (PAM) per plaque compared to R-;AD+ mice (p < 0.001; Fig. 2D). In addition, there was 2-fold more phagocytosis of Aβ_42_ protein per PAM in R+;AD+ microglia compared to R-;AD-microglia (p < 0.0001; Fig. 3A). Endolysosomal trafficking mediated by the myeloid immune receptors, CLEC7A and TREM2 was also enhanced in ACE expressing microglia. CLEC7A (also known as DECTIN1) is a c-type lectin receptor upregulated in microglia in 5xFAD mice and during neurodegeneration (44–46). Real-time fluorescence measurement in primary microglia isolated from R+;AD- and R-;AD- mice treated with Zymosan conjugated to pHrodo, which binds and activates CLEC7A receptors (Fig 3B), showed significantly increased phagocytosis and endolysosome loading (Fig 3C), and degradation (Fig. 3D) in ACE-expressing microglia. Similarly, TREM2 is up-regulated in DAM in 5xFAD mice (44, 47) and in microglia from patients with AD (30). TREM2 binds Aβ_42_ to activate microglial immune response and function, including protein clearance and cytokine production (47). Real-time fluorescence measurement of microglia treated with oligomeric Aβ_42_ (oAβ_42_) conjugated to pHrodo (Fig. 3E) also showed significantly increased phagocytosis and lysosome loading (Fig 3F), and increased degradation (Fig 3G) in microglia isolated from R+;AD- compared to R-;AD- mice. Together, these results suggest that ACE expression in microglia enhances phagocytosis, endolysomal trafficking and degradation of extracellular Aβ through mechanisms that involve TREM2 and CLEC7A receptor binding and downstream signaling (31). Increased Aβ_42_ trafficking was associated with an increase in the volume of the CD68+ endolysosomal compartment. The endolysosomal compartment volume was expanded by 5.4- fold in R-;AD+ compared to R-;AD- microglia (p < 0.0001), and it was enhanced another 1.7-fold with ACE expression in microglia (R+;AD+) compared to R-;AD+ microglia (p < 0.01, Fig. 3H). These results suggest that decreased amyloid plaque burden in the brain of R+;AD+ mice is at least in part due to increased recruitment of microglia to Aβ-containing plaques and to their enhanced phagocytosis and endolysosomal degradation of Aβ_42_ within them.

**Fig. 3.**
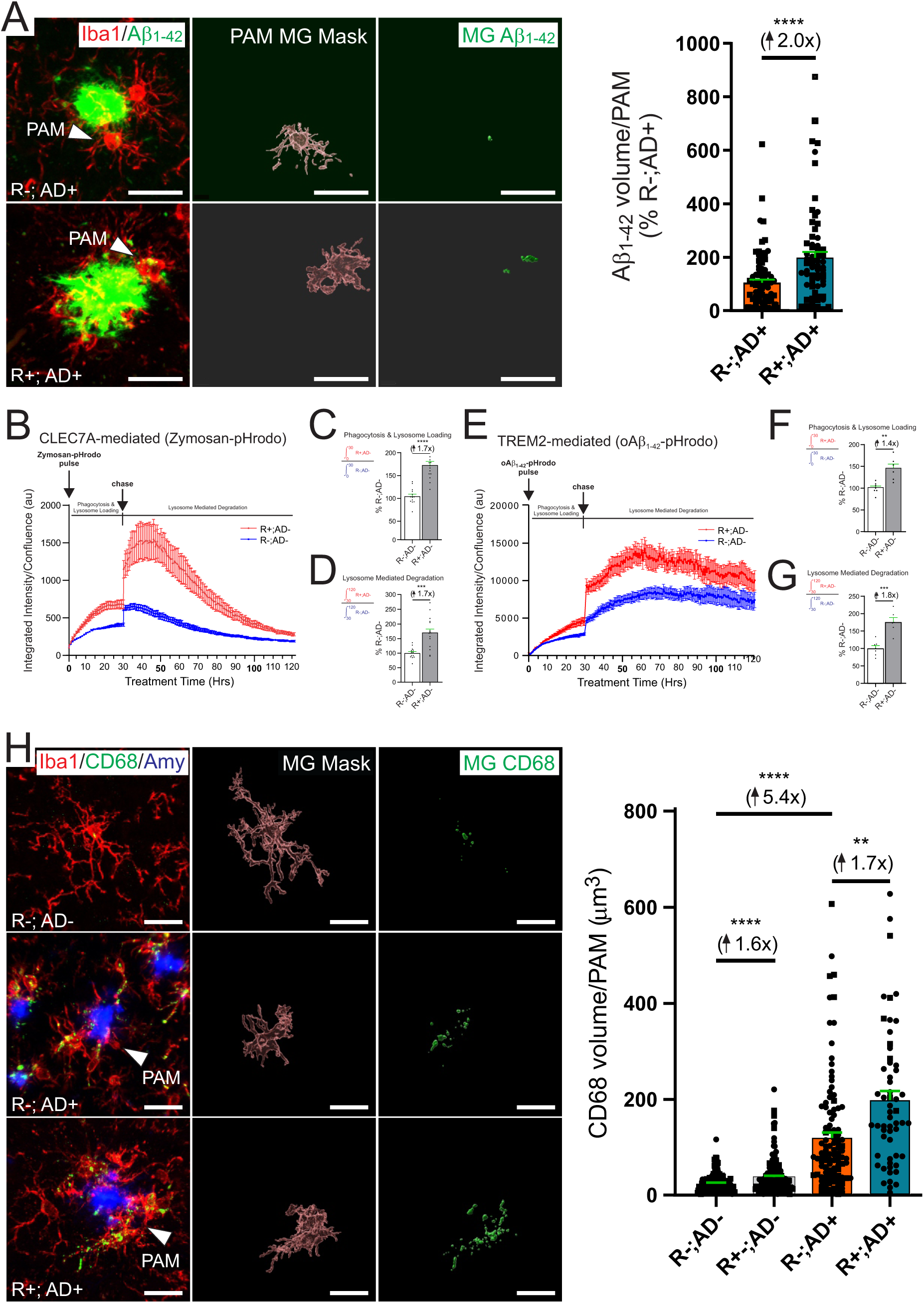
Increased Aβ phagocytosis and endolysosomal trafficking in ACE expressing microglia. **(A)** Aβ protein phagocytosis was measured using Iba1 immunofluorescence to mask individual PAM in 3-dimensions on 50 µM thick tissue sections with iMaris image analysis software to determine the volume of Aβ_1-42_ within them. Microglia from R+;AD+ mice had 2-fold more internalized Aβ_1-42_ compared to R-;AD+ microglia (p < 0.0001). **(B)** Primary microglia isolated from R+;AD- (R+) and R-;AD- (R-) mice were grown in culture and treated with Zymosan conjugated to pHrodo (Zymosan-pHrodo) which binds to CLEC7A receptors or **(E)** oligomeric Aβ_1-42_ (oAβ_1-42_) conjugated to pHrodo (oAβ-pHrodo) which binds to Trem2 receptors. The molecules fluoresce when internalized and trafficked to the acidified endolysomal compartment. Cell fluorescence intensity was measured by Incucyte for 120 hours to identify engulfment and loading of pHrodo into microglia from the culture media (pulse) and then endolysosomal trafficking and degradation was measured when it was washed out from the media (chase) for R+;AD- (red) and R-;AD- (blue) microglia. **(C)** Phagocytosis and early lysosome loading of Zymosan-pHrodo (0-30 hours) was increased by 1.7-fold (p < 0.0001) and **(D)** lysosomal trafficking and degradation (30-120 hours) was increased by 1.7-fold (p < 0.05) compared to R-;AD- microglia. Similarly, **(F)** oAβ-pHrodo phagocytosis and early lysosome loading was increase 1.4-fold (p < 0.01) and **(G)** lysosomal trafficking and degradation was increased by 1.8- fold (p < 0.001) in ACE expressing (R+) microglia. **(H)** Immunofluorescence to detect Iba1+ microglia (red), CD68+ endolysomal compartment (green), and Amylo-Glo to detect amyloid plaques (blue) showed increased endolysosomal trafficking in R-;AD+ and R+;AD+ compared to R-;AD- PAM. Imaris filament tracing was used to generate 3D masks from 50 µM thick tissue sections representing individual PAM and then to quantify the volume of CD68 protein within them. R-;AD+ PAM had 5.4-fold increase in CD68 labeling compared to R-;AD- microglia, reflecting increased endolysosomal trafficking within PAM surrounding amyloid plaques (p < 0.0001). In R+;AD+ PAM, there was an additional 1.7-fold increase in CD68 labeling compared to R-;AD+ PAM (p < 0.01), suggesting that ACE expression in microglia enhances endolysosomal trafficking and protein degradation within them in vivo. (A, H scale bar = 20 µM; square symbol = male and circular symbol = female)

### ACE expression in microglia reduces neuron and synapse degeneration in 5xFAD mice

By 9-months of age, neuron and synapse degeneration that is correlated with abnormalities in learning and memory behavior are present in 5xFAD mice (48). Consistent with published reports, R-;AD+ mice had significant neuron loss in previously identified vulnerable regions of the brain (49), with a 1.7-fold decrease in hippocampal subicular neuron density (p < 0.0001) and a 1.3-fold decrease in layer V forebrain cortical neuron density (p < 0.01). Interestingly, ACE expression in microglia significantly reduced neuron degeneration and restored neuron density to control (R-;AD-) levels in the subiculum and in layer V cortical forebrain neurons (Fig 4A).

**Fig. 4.**
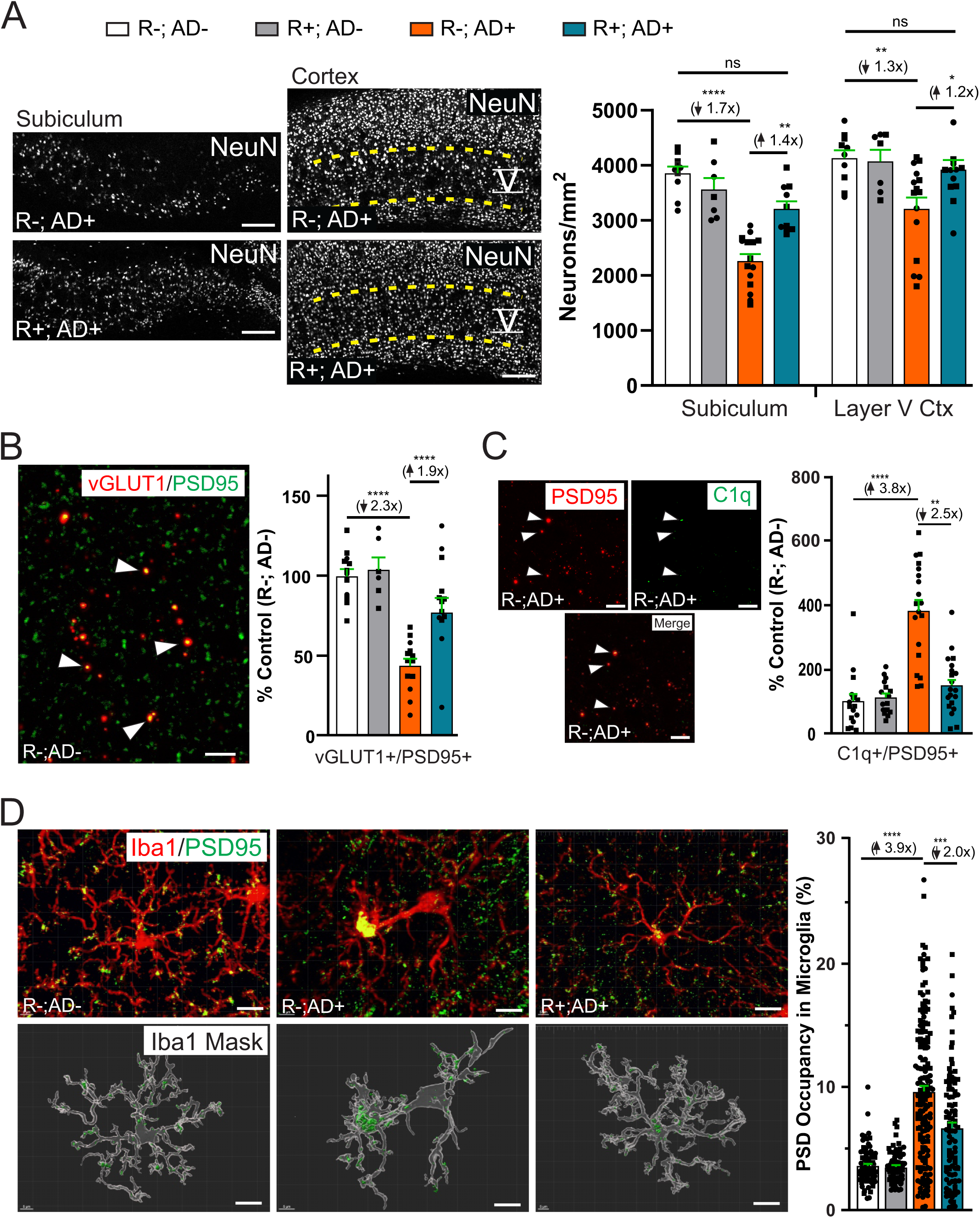
ACE expression in microglia reduces neuron and synapse degeneration in 5xFAD mice. **(A)** Neurons in the subiculum of the hippocampal formation (Sub) and layer V forebrain cortical neurons (V) were labeled by immunofluorescence with an antibody to NeuN, and neuron density in these regions was measured (representative images from R-;AD+ and R+;AD+ shown). Neuron density was decreased by 1.7-fold in Sub (p < 0.0001) and by 1.3-fold in V (p < 0.01) in R-;AD+ compared to R-;AD- mice, consistent with neuron loss in these brain regions in 5xFAD mice. Neuron density in R+;AD+ Sub and V were increased to levels found in R-;AD- mice. **(B)** Synaptic density in the forebrain was measured using immunofluorescence to detect presynaptic excitatory (vGlut1) and postsynaptic (PSD95) specializations. Excitatory synapses with co-localized vGlut1 and PSD95 (arrowheads) were decreased by 2.3-fold in R-;AD+ compared to R-;AD- mice (p < 0.0001) and they were increased by 1.9-fold (p < 0.0001) in R+;AD+ compared to R-;AD- mice. **(C)** PSD95-labeled postsynaptic specializations were co- labeled with C1q 3.8-fold more frequently in R-;AD+ compared to R-;AD- mice (p < 0.0001), consistent with increased tagging and synapse loss that occurs in AD+ brains. C1q labeling was reduced by 2.5-fold in R+;AD+ compared to R-;AD+ mice (p < 0.01), consistent with a decrease of synapse loss when ACE is expressed in microglia (PSD95, red; C1q green). **(D)** 3D reconstruction of individual CA1 microglia from 50 µM thick tissue sections were masked to quantify the volume of internalized PSD95 within them. Postsynaptic PSD95 phagocytosis (green) was increased 3.9-fold in Iba1 labeled microglia (red) from R-;AD+ compared to R-;AD- mice (p < 0.0001). Phagocytosis of PSD95 was reduced by 2-fold in R+;AD+ compared to R-;AD+ mice (p < 0.001), consistent with reduced synapse loss when ACE is expressed in microglia. (A, scale bar = 200 µM; B, C, scale bar = 1 µM; D, scale bar = 10µM; male mice = round symbol and female mice = square symbol).

Excitatory synapse loss is correlated with neuron loss from excitatory glutaminergic neurons in the forebrain and the loss is reduced by ACE expression in microglia. Excitatory synapses marked by puncta labeled by the excitatory presynaptic specialization marker, vGlut1 and co- localized with the postsynaptic specialization marker, PSD95 were reduced by 2.3-fold in R-;AD+ compared to R-;AD- mice (p < 0.0001). ACE expression in microglia increased excitatory synapse density by 1.9-fold (p < 0.0001) to near control (R-;AD-) levels (Fig. 4B). Synapses are eliminated, at least in part, by mechanisms that involve labeling them with the complement component C1q to “mark” them for microglia-dependent elimination (50). Consistent with the loss of synapses in the forebrain of R-;AD+ mice, PSD95 labeled synapses were 3.8-fold more frequently labeled with C1q in R-;AD+ compared to R-;AD- mice (p < 0.0001). Moreover, in R+;AD+ mice, where excitatory synapses were increased compared to R-;AD+ mice, there was a 2.5-fold decrease in the frequency of C1q labeling in synapses marked by PSD95 (p < 0.01; Fig. 4C). Similarly, in R-;AD+ mice, the decreased density of PSD95 postsynaptic specializations was accompanied by a 3.9-fold increase in phagocytosis of PSD95 by microglia (p < 0.0001) which was decreased by 2.0-fold in microglia from R+;AD+ compared to R-;AD+ mice (p < 0.001; Fig. 4D). Taken together, these results suggest that ACE expression in microglia reduces neuron and synapse degeneration in 5xFAD mice, at least partly through decreasing PSD95 phagocytosis and decreasing C1q-mediated synapse turnover by microglia.

### ACE expression in microglia restores learning and memory abnormalities in 5xFAD mice

5xFAD mice have learning and memory abnormalities that correlate with amyloid plaque deposition and degeneration of neurons and synapses (51). Mice of all genotypes (R-;AD-, R+;AD-; R-;AD+ and R+;AD+) had similar body weights (Fig. S6A), locomotor behavior (Fig. S6B, C) and anxiety-like behavior in open field testing (Fig. S6D, E). However, in a hippocampus-dependent Barnes maze memory test, 9 month old R-;AD+ mice showed significant learning abnormalities with increased time to locate an object (latency) after 5 days of training compared to R-;AD- mice (p < 0.0001). With ACE expression in microglia, R+;AD+ mice had significantly decreased learning latency compared to R-;AD+ mice over a 5 day training period (p < 0.001; Fig. 5A). In a memory task performed two-days after training to locate the object, R-AD+ mice had memory impairment with a 3.9-fold increase in latency compared to R-;AD- mice (p < 0.0001). Interestingly, latency was significantly reduced by 1.6- fold in R+;AD+ compared to R-;AD+ mice (p < 0.001; Fig. 5B). Similar results were obtained when mice were trained to find a new object location (reversal memory). Latency to locate the new object in R-;AD+ mice was significantly increased 3.5-fold compared to R-;AD- mice (p < 0.0001) and it was significantly decreased by 1.3-fold (p < 0.05) in R+;AD+ compared to R-;AD+ mice (Fig. 5C).

**Fig. 5.**
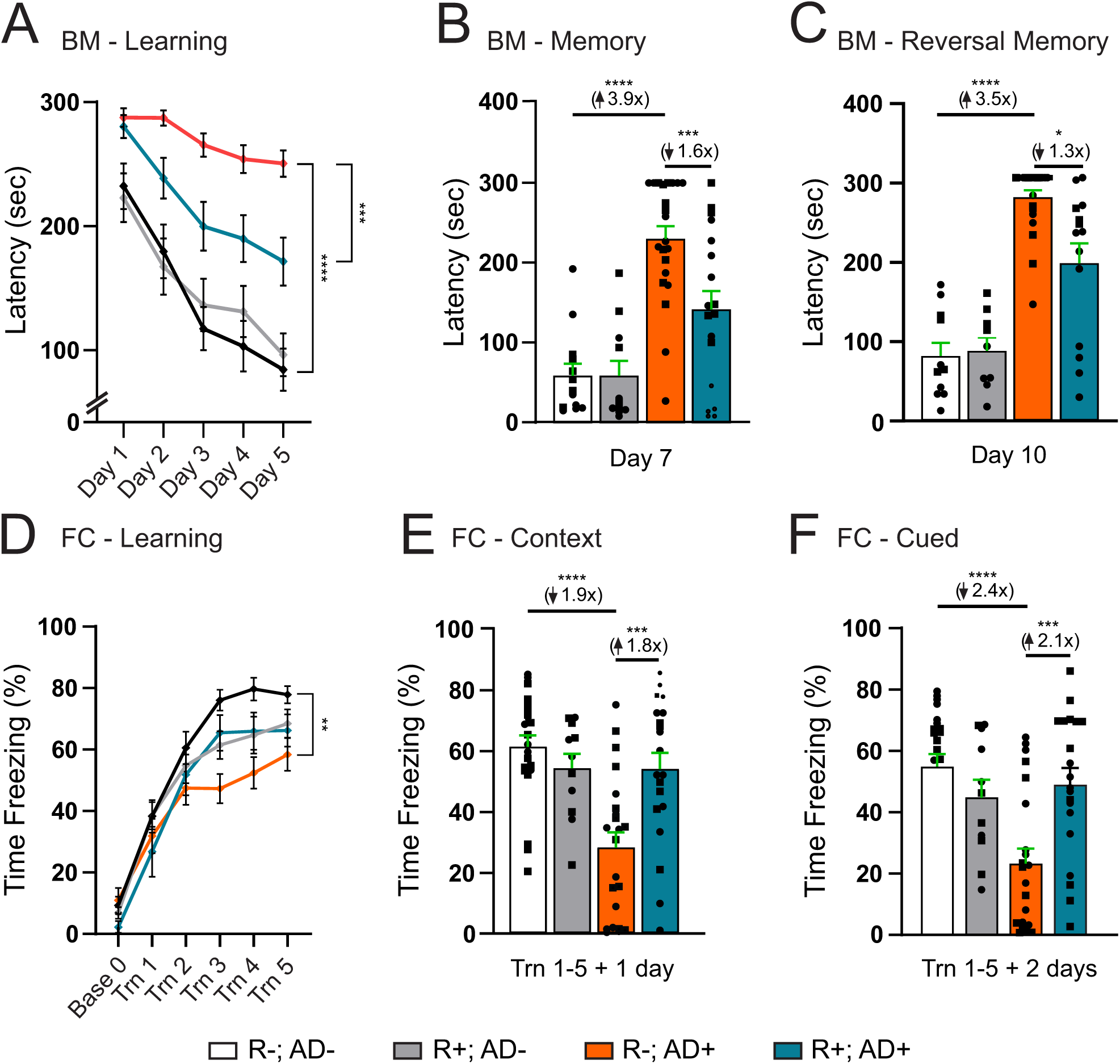
ACE expression in microglia significantly improves learning and memory abnormalities in 5xFAD mice. Barnes maze and fear conditioning were used to test hippocampal-dependent and amygdala-dependent learning and memory behavior. **(A)** Training for five days to locate an object in a Barnes maze showed significant increase in time to locate the object (latency) in R-;AD+ compared to R-;AD- mice (p < 0.0001). However, latency to learn the location of the object was significantly reduced in R+;AD+ compared to R-;AD+ mice (p < 0.001). **(B)** In a memory test, latency to locate the object was significantly increased in R-;AD+ by 3.9-fold compared to R-;AD- mice two days after training (p < 0.0001) and latency to locate the object was significantly decreased in R+;AD+ by 1.6-fold compared to R-;AD+ mice (p < 0.001). **(C)** In a reversal memory test, by moving the object to a new location and re-training, R-;AD+ mice again had a significant 3.5-fold increased latency to locate the object compared to R-;AD- mice (p < 0.0001), which was significantly decreased by 1.3-fold in R+;AD+ compared to R-;AD+ mice (p < 0.05). **(D)** In fear conditioning behavioral tests, R-;AD+ mice had significantly impaired learning of contextual and cued stimuli during training compared to R-;AD- mice (p < 0.01). **(E)** In contextual memory testing 1 day after training, R-;AD+ mice had significant memory impairment with a 1.9-fold decrease in freezing time compared to R-;AD- mice (p < 0.0001). Memory was significantly improved with freezing time increased by 1.8-fold in R+;AD+ compared to R-;AD+ mice (p < 0.001). **(F)** In cued memory testing 2 days after training, R-;AD+ mice had significant memory impairment with a 2.4-fold decrease in freezing time compared to R-;AD- mice (p < 0.0001). Memory was significantly improved with a 2.1-fold increase in freezing time in R+;AD+ compared to R-;AD+ mice (p < 0.001).

Freezing time in a fear conditioning assay is increased when an animal perceives an adverse conditioned stimulus that is either environmentally contextual (hippocampal-dependent) or auditory cued (amygdala-dependent) when paired with an unconditioned shock stimulus. Time freezing was significantly decreased during 5 training sessions in R-;AD+ mice compared to R-;AD- mice (p < 0.01), and it was similar between all other genotypes (Fig. 5D). However, in a contextual memory test performed 1 day after training, R-;AD+ mice had a significant 1.9-fold reduction in time freezing compared to R-;AD- mice (p < 0.0001), indicating their impaired memory of the previous conditioning. ACE expression in microglia in R+;AD+ mice significantly increased freezing time by 1.8-fold in R+;AD+ compared to R-;AD+ mice (p < 0.001; Fig. 5E), indicating improved memory of the previous conditioning. Similarly, in an auditory cued memory test performed 2 days after training, R-;AD+ mice had a significant 2.4-fold decrease in time freezing (p < 0.0001) compared to R-;AD- mice, which was significantly increased by 2.1-fold in R+;AD+ compared to R-;AD+ mice (p < 0.001). Thus, in addition to reducing Aβ plaque load and rescuing neuron and synapse loss, ACE expression in microglia significantly reduces learning and memory abnormalities in 5xFAD mice.

### ACE expression in microglia drives molecular signaling pathways mediating endolysosomal trafficking, Syk tyrosine kinase signaling and cell migration

To characterize the molecular changes induced by ACE in microglia, single nuclei were isolated from 6-month-old R-;AD-, R+;AD-, R-;AD+ and R+;AD+ frozen forebrains. Microglia nuclei were enriched by fluorescence activated cell sorting (FACS) to diminish NeuN+ neuron, Sox10+ oligodendroglia, and Sox9+ astrocyte nuclei in the samples (Fig. S7A) before single nuclear capture and reverse transcription (10x Genomics) (Fig. S7B). snRNAseq and UMAP analysis identified 10 transcriptionally distinct cell clusters (Fig. S7C) that were assigned to their cellular identity by mapping well-established lineage associated gene expression to them (Fig. S7D). Microglia were enriched approximately 3-5-fold and accounted for 10% of all nuclei sequenced (Fig. S7C). Microglia were represented by two unique cell clusters that expressed the microglia lineage-enriched markers, Csf1r, Cx3cr1, Tmem119, C1qa, Hexb and P2ry12 (Fig. S7D, E).

The two microglia clusters were pooled to perform a secondary UMAP analysis to identify 8 transcriptionally distinct microglia sub-clusters that were further demultiplexed to compare genotype-dependent gene expression and cell distribution within them (Fig. 6A). Gene expression analysis identified microglia sub-clusters with gene expression profiles similar to previously reported homeostatic microglia (HM; sub-cluster 1), transitional microglia (TM; sub- cluster 2) and disease-associated microglia (DAM) (sub-cluster 3, designated as DAM-A) (Fig. S8) (44). In addition, a previously uncharacterized small DAM sub-cluster was identified (sub- cluster 5) and designated as DAM-B (Fig. 6B). The frequency of microglia in DAM-A expanded from 0.3% to 26.6% (χ^2^ test, p < 10^-34^) and in DAM-B, microglia expanded from 2.8% to 11.4% (χ^2^ test, p < 10^-9^) in R-;AD+ compared to R-;AD- mice. With ACE expression in microglia, the frequency of DAM-A was increased further by 4.6% and in DAM-B it was decreased by 4.9% in R+;AD+ compared to R-;AD+ mice (Fig. 6A). Thus, ACE expression in microglia shifts transcriptionally distinct DAMs by increasing DAM-A and decreasing DAM-B frequency.

**Fig. 6.**
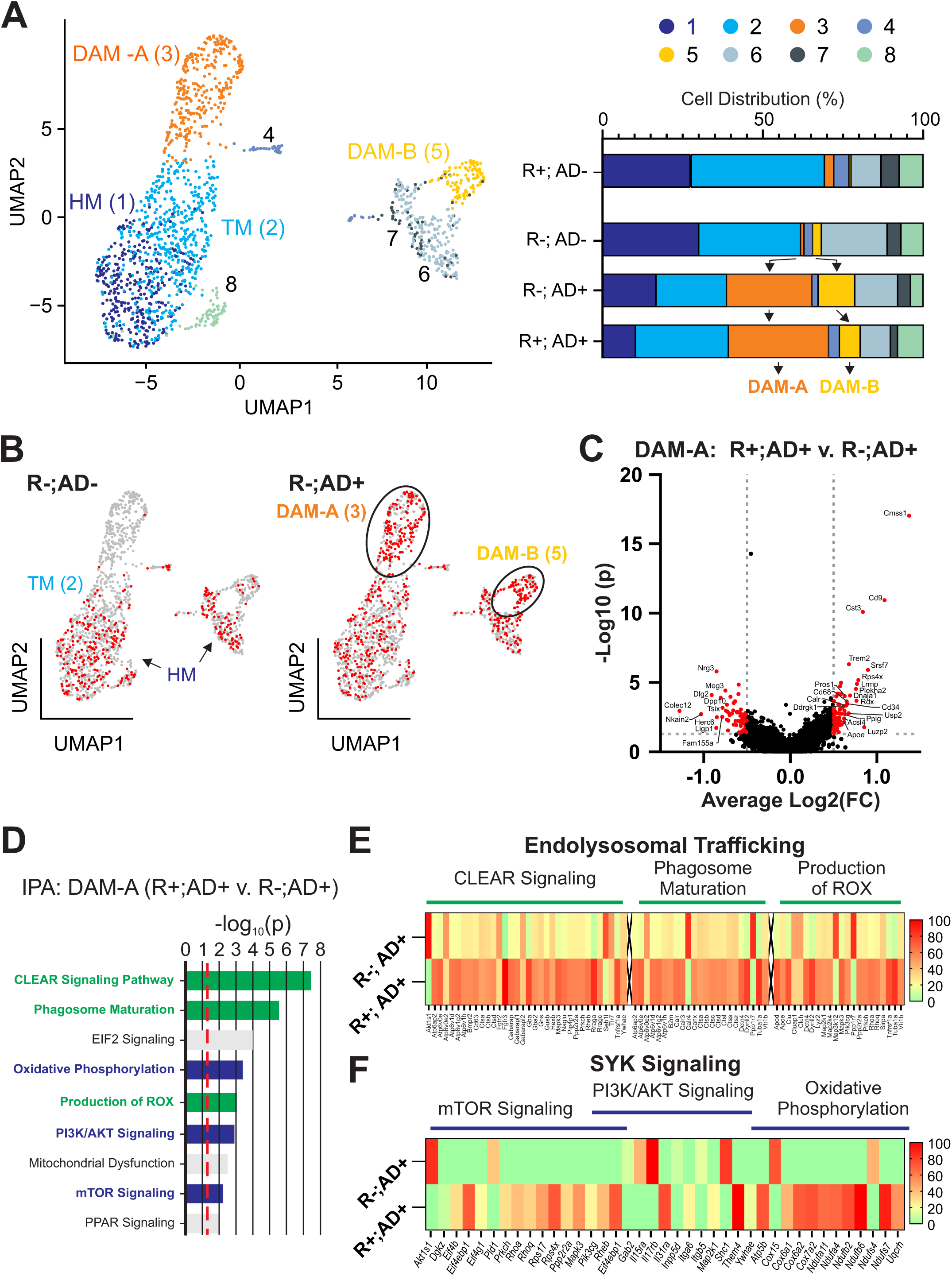
ACE expression in disease associated microglia (DAM) modulates endolysosomal trafficking and SYK signaling. **(A)** snRNAseq and UMAP subcluster analysis of microglia from the forebrain of 9-month-old R-;AD-, R+;AD-, R-;AD+ and R+;AD+ mice identified 8 transcriptionally distinct microglia subclusters. Several microglia subclusters had gene expression profiles consistent with previously described homeostatic (HM; subcluster 1), transitional (TM; subcluster 2) and disease associated (DAM-A; subcluster 3) microglia. The DAM-A subcluster contained previously identified TREM2-independent DAM stage 1 (DAM-1) and TREM2-dependent DAM stage 2 (DAM-2) microglia (44). A smaller and poorly defined DAM subcluster (subcluster 5) was identified and designated as DAM-B. The frequency of DAM-A and DAM-B were highly altered by genotype. The frequency of DAM-A was increased from 0.3% to 26.6% and in DAM-B they were increased from 2.8% to 11.4% in R-;AD+ compared to R-;AD- mice. With ACE expression in microglia the frequency of DAM-A was increased by 4.6% (to 31.2%) and in DAM-B they were decreased by 4.9% (to 6.5%). **(B)** DAM- A and DAM-B microglia were only observed in the presence of amyloid in AD+ brains, defining them as DAM, and their frequency was skewed by ACE expression in them. **(C)** Ingenuity Pathway Analysis (IPA) identified many highly significantly altered signaling pathways in DAM-A R+;AD+ compared to R-;AD+ microglia. Although human ACE (hACE) was elevated in R+ microglia, it was not detected in the mouse sequencing data that used the mouse reference genome for sequence mapping. **(D)** Endolysosomal trafficking involving CLEAR signaling, phagosome maturation and production of ROX as well as SYK signaling involving mTOR, PI3/AKT signaling, and oxidative phosphorylation were upregulated by ACE expression in microglia. **(E, F)** IPA analysis identified pathways associated with endolysosomal trafficking and SYK signaling that where upregulated by ACE expression in DAM-A (R+;AD+).

The majority of DAM were represented by DAM-A which contained both TREM2-independent DAM stage 1 (DAM-1) and TREM2-dependent DAM stage 2 (DAM-2) that were previously described in 5xFAD mice (Fig. S8) (52). ACE expression in DAM-A significantly modulated the expression of 165 genes, including the up-regulation of two late onset AD associated genes, Apoe and Trem2 (8, 53), up-regulation of 15 additional previously described DAM genes (such as, CD9, Cstd and Ctsl) and down-regulation of 3 previously described DAM genes (Colec12, Dkk2 and Myo5a) (44) (Fig. 6C, Table S1). In addition, CD68 which was also upregulated at the protein level in the expanded endolysosomal compartment of R+;AD+ microglia (Fig. 3H), was also upregulated at the transcriptional level by snRNAseq (Fig. 6C).

To better understand how ACE expression in microglia modulates their function to decrease Aβ plaque load, neuron/synapse degeneration and learning and memory impairments, Ingenuity Pathway Analysis (IPA) was performed on the genes differentially expressed between R+;AD+ and R-;AD+ DAM-A. Gene regulatory pathways related to endolysosomal trafficking and spleen tyrosine kinase (SYK) signaling were significantly modulated by ACE expression in DAM-A (Fig. 6D, Table S2). Increased endolysosomal trafficking was characterized by significantly altered gene expression in pathways mediating CLEAR signaling, phagosome maturation and reactive oxygen species (ROX) production (Fig. 6E, Table S3). CLEAR signaling has a major role in lysosomal maturation and cellular clearance which recruits phagosome maturation mechanisms and ROX production (54, 55). These molecular changes are consistent with increased internalization of Aβ_42_ and the expanded CD68+ endolysosomal compartment that was observed *in vivo* in ACE-expressing PAM from R+;AD+ compared to R-;AD+ mice (Fig. 3A, H).

Recent studies indicate that CLEC7A and TREM2 mediate microglia activation in response to Aβ binding through phosphorylation of SYK tyrosine kinase which in turn activates downstream signaling pathways such as mTOR and PI3K/AKT, and functional pathways that regulate oxidative phosphorylation and cell migration (30, 32, 56). Interestingly, IPA identified many downstream SYK signaling pathways that were highly upregulated in R+;AD+ compared to R-;AD+ DAM-A, such as mTOR and PI3K/AKT signaling (Fig 6F, Table S4). Similarly, genes involved in oxidative phosphorylation were generally upregulated, which is consistent with a previously identified role for ACE in enhancing immune function at least in part through increasing oxidative phosphorylation and ATP biogenesis in other myeloid-derived cells, such as macrophages and neutrophils (57). Similarly, the IPA function for *Movement of Cells* was highly modulated by ACE expression in microglia (Fig S9, Table S5), which likely correlates with increased migration of microglia that accumulate around plaques in R+;AD+ relative to R-;AD+ mice *in vivo* (Fig. 2D).

To better understand the role of ACE-expression in DAM-A and their spatial relationship to Aβ plaques in the brain, we generated tissue microarrays from the hippocampus of 9-month-old male and female R-;AD-, R+;AD-, R-;AD+ and R+;AD+ mice, and examined them using whole transcriptome analysis with the Nanostring GeoMx digital spatial profiling (DSP) platform. Morphology markers for Iba1 to identify microglia, Aβ_42_ to identify plaques and Syto13 to identify cell nuclei were used to guide the selection of regions of interest (ROI) containing plaque-associated microglia (PAM) and non-plaque-associated microglia (nPAM) for gene expression analysis (Fig 7A). The Iba1 fluorescence channel was used to generate ultraviolet light masks to elute probes bound to RNA within microglia in the ROI (Fig. 7B). In previous studies, DAM appeared to correspond to PAM based on analysis of a small number of DAM proteins that were upregulated primarily in PAM relative to nPAM (44). DSP analysis identified many genes that were deregulated in PAM compared to nPAM (Fig. S10A, Table S6) and there was a highly significant correlation between genes upregulated in PAM by DSP with those upregulated in DAM-A by snRNAseq (p < 0.0001) (Fig. S10B). Thus, DAM-A appear to primarily represent PAM, while DAM-B represent a small population of DAM that are not apparently associated with plaques.

**Fig. 7.**
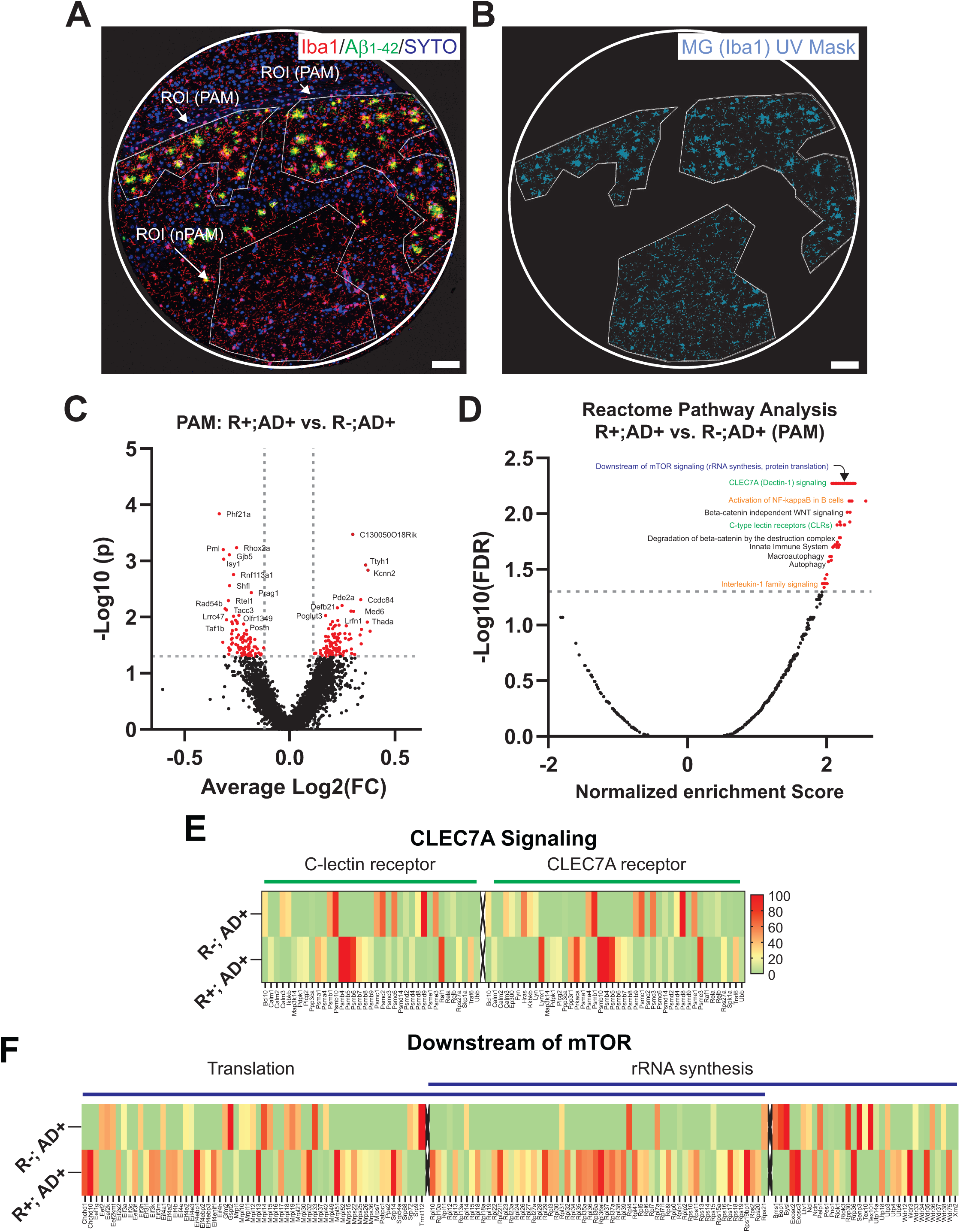
ACE modulates CLEC7A, mTOR and inflammatory signaling in microglia. **(A)** Regions of interest (ROI) containing plaque associated microglia (PAM) and non-plaque associated microglia (nPAM) were selected using immunofluorescence morphology markers to localize microglia (Iba1) and plaques (Aβ_1-42_), and the nuclear stain SYTO Green. **(B)** Nanostring digital spatial profiling (GeoMx) and whole transcriptome analysis were used on regions of interest (ROI) selected using the morphology markers, and the Iba1 channel was used to generate a UV mask to photo-elute probes hybridized primarily to microglia. **(C)** ACE expression in PAM significantly modulated gene expression within them. Although human ACE (hACE) was highly expressed in microglia, it was not detected by the mouse-specific probes in GeoMx. **(D)** Reactome pathway analysis showed that many of the genes regulated by ACE in PAM modulate signaling pathways related to mTOR, CLEC7A, and inflammatory signaling. **(E)** ACE expression in PAM modulates many genes related to the c-lectin receptor CLEC7A signaling in microglia and **(F)** genes related to signaling pathways downstream of mTOR. (A, B scale bar = 100 µM)

ACE expression in microglia led to significant differential gene expression in R+;AD+ compared to R-;AD+ PAM (Fig. 7C, Table S7), although the sensitivity to detect differentially expressed genes was probe dependent and appeared to be decreased compared to snRNAseq analysis performed on DAM-A (Fig. 6C). Nevertheless, Reactome Pathway Analysis (RPA) highlighted the functional impact of ACE expression in PAM (Fig. 7D, Table S8) and it showed similarity to pathways identified by IPA performed on gene expression identified by snRNAseq on DAM-A. The pathways most significantly modulated by ACE in PAM were related to C-type lectin receptor and CLEC7A receptor signaling (Fig. 7E, Table S9), pathways consistent with SYK signaling downstream of mTOR, which included protein translation and rRNA synthesis (Fig. 7F, Table S10), and pro-inflammatory signaling through NFκB and IL1-β (Fig. S11, Table S11). Taken together, these results indicate that ACE expression in microglia modulates their ability to process Aβ_42_ by driving signaling through TREM2 and CLEC7A receptors, and their downstream tyrosine kinase effector SYK, to potentiate many recently identified functions of SYK signaling in microglia, including increased plaque association, Aβ engulfment, neuroprotection, and decreased Aβ plaque formation (30–32, 58).

Activated intracellular SYK signaling can be identified by its phosphorylation at Y352 (pSYK) (59, 60). There was no detectable difference in the amount of pSYK in microglia mediated by ACE expression between R+;AD+ and R-;AD+ nPAM (Fig. 8A). However, pSYK was increased by 2.0-fold (p < 0.05) in PAM compared to nPAM in R-;AD+ mice (Fig. 8A), consistent with previous results indicating that SYK signaling is increased in PAM in AD (30, 58). Interestingly, in ACE expressing PAM from R+;AD+ mice, there was a further 2.4-fold increase in the amount of pSYK within them compared to R-;AD+ PAM (p < 0.0001), consistent with the pathway analyses showing molecular signatures related to increased SYK signaling in ACE expressing microglia (Fig. 8A). To test whether increased Aβ_42_ uptake and endolysosomal trafficking in ACE expressing microglia is mediated by increased SYK signaling (Fig. 3), primary microglia were isolated from R-;AD- and R+;AD- mice and cultured in vitro. Treatment of ACE expressing microglia from R+;AD- mice with oligomeric Aβ_42_ (oAβ_42_) showed a 1.7-fold increase in uptake and trafficking into the CD68 endolysosomal compartment when compared to R-;AD- microglia (p < 0.0001) (Fig. 8B), consistent with a role for ACE in potentiating oAβ_42_ uptake and trafficking that was previously shown to be mediated by SYK signaling (30–32). Accordingly, treatment with the selective SYK inhibitor PRT062607 (P505) decreased oAβ_42_ uptake and endolysosomal trafficking by 1.7-fold in the absence of ACE expression in R-;AD- microglia (p < 0.01), and it decreased uptake and degradation of oAβ_42_ by 2.2-fold in ACE expressing microglia from R+;AD- mice (p < 0.0001). Similar results were obtained when gene silencing was used to specifically knockdown SYK in primary microglia (Fig. 8C). Reduction of Syk expression in R-;AD- microglia treated with RNAi (*i*) reduced oAβ_42_ uptake and trafficking by 2.3-fold compared to RNAscramble (*SCR*) treated microglia (p < 0.05). In R+;AD- microglia treated with *SCR*, oAβ_42_ uptake and endolysosomal trafficking was increased by 1.8-fold compared to R-;AD- *SCR* treated microglia (p < 0.001), and in R+;AD- microglia, *i* treatment reduced oAβ_42_ uptake and trafficking by 2.7-fold compared to *SCR* treated cells (p < 0.0001). SYK signaling abrogated by either pharmacologic inhibition or knockdown, led to reduction of the potentiating effects of ACE expression in R+;AD- microglia on oAβ_42_ uptake and trafficking to levels statistically indistinguishable from non-ACE expressing microglia. Taken together, these results indicate that ACE expression in microglia potentiates oAβ_42_ uptake and endolysosomal trafficking through SYK-dependent mechanisms that are known to coordinate cellular processes mediating Aβ engulfment, plaque association, decreased plaque formation and neuroprotection (30–32).

**Fig. 8:**
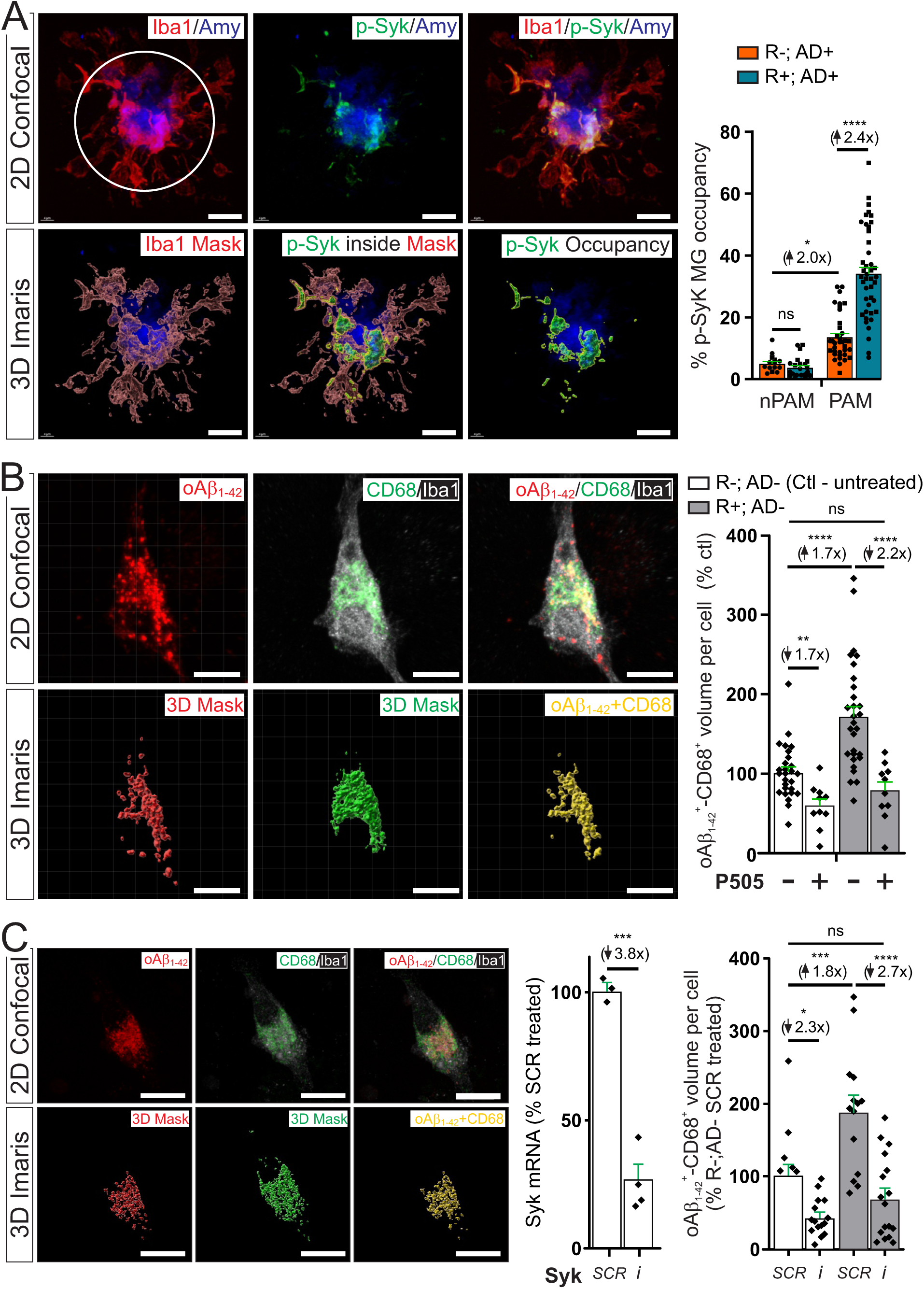
ACE expression in microglia enhances SYK phosphorylation and SYK-dependent Aβ phagocytosis and endolysosomal trafficking. **(A)** SYK phosphorylation was increased 2-fold in PAM relative to nPAM (p < 0.05). With ACE expression in microglia, levels of p-SYK within PAM increased by 2.4-fold (p < 0.0001), consistent with increased SYK signaling relative to microglia not expressing ACE. (circle denotes the region within 10 µM of a plaque edge used to classify microglia as PAM if cell body or process encroach within the circle). **(B)** Primary microglia treatment with oAβ_42_ resulted in a 1.7-fold increase in oAβ_42_ phagocytosis and trafficking to the CD68+ endolysosomal compartment when ACE was expressed (R+; AD-) compared to microglia without ACE expression (R-; AD-) (p < 0.0001). Treatment of microglia with the SYK inhibitor (P505 - PRT062607) decreased oAβ_42_ phagocytosis and trafficking by 1.7-fold in microglia not expressing ACE (p < 0.0001) and abrogated trafficking by 2.2-fold (p < 0.0001) in microglia expressing ACE. **(C)** RNAi was used to decrease SYK expression in primary microglia (efficacy of SYK knockdown shown using RT-PCR). RNAi (*i*) treatment of R-;AD- microglia reduced oAβ_42_ phagocytosis and trafficking by 2.3-fold compared to scramble (*SCR*) treated R-;AD- microglia. While microglia expressing ACE (R+;AD-) treated with *SCR* showed a 1.8-fold increase in oAβ_42_ phagocytosis and trafficking compared to *SCR* treated R-;AD- microglia, oAβ_42_ phagocytosis and trafficking was reduced by 2.7-fold in *i* treated R+;AD- microglia compared to *SCR* treated R+;AD- microglia. (A-C, scale bar = 10 µM).

## DISCUSSION

ACE haploinsufficiency or administration of a CNS penetrant ACE catalytic activity inhibitor exacerbates amyloid deposition and neurodegeneration in hAPPSw mice (26). In humans, increased cortical ACE expression protects against development of AD and the effect is not associated with blood pressure regulation (61). Previous studies showed that ACE expression in myeloid-derived macrophages and neutrophils potentiates their immunologic function (27, 28, 62) and that ACE expression in peripheral macrophages in ACE^10/10^ mice reduces Aβ deposition, neurodegeneration and improves learning and memory deficits in APP/PS1 amyloid producing mice. However, we found that ACE^10/10^ mice also have high expression of ACE in microglia which raised the possibility that ACE activity in microglia, rather than in peripheral macrophages was the primary reason reduced Aβ deposition and neuroprotection (63) (Fig. S1). Here, we selectively expressed ACE in microglia/CAMs and found that it potentiates the ability of microglia to process Aβ. We found that in R+;AD+ mice there was decreased Aβ plaque burden, reduced neurodegeneration and excitatory synapse loss, and reduced learning and memory impairments. Moreover, ACE-expressing microglia in R+;AD+ mice were recruited to plaques in higher number, and they engulfed more Aβ that was targeted to an expanded endolysosomal compartment. Accordingly, pathway analysis from snRNAseq and DSP studies identified many ACE-regulated pathways in DAM. For example, there was evidence of altered pro-inflammatory NFκB and IL-1β signaling, and significant alteration of the IPA *Movement of Cells Network* in DAM, which may drive their altered immune activation and increased aggregation around Aβ plaques. Similarly, while in vitro and in vivo labeling of the endolysosomal compartment with CD68 showed increased endolysosomal trafficking of Aβ in ACE expressing microglia, functional pathway analysis also showed upregulation of the *CLEAR Signaling* network, a master regulator of lysosomal biogenesis and dynamics, and *Phagosome Maturation* and *Production of Reactive Oxygen Species*. Specifically, expression of several pH-sensitive cathepsin lysosomal proteases (cathepsin A, B, C, D, L, S and Z) were markedly upregulated in ACE-expressing DAM, which may have contributed to their enhanced ability to degrade Aβ. This appeared to be a cell-autonomous enhancement of microglial function as ACE-expressing microglia derived from R+;AD- mice treated in vitro with Zymosan and oAβ_42_ conjugated to pHrodo showed clear evidence of enhanced phagocytosis and Aβ degradation mediated by their receptors, CLEC7A and TREM2, respectively.

The complement component C1q opsonizes damaged synaptic membranes, and in AD models with Aβ deposition there is increased C1q-labeling of synapses that leads to widespread synapse loss mediated by microglia (50). We found that while C1q subunits, C1qa, C1qb and C1qc were upregulated in ACE-expressing DAM, PSD95+ post-synaptic membranes were less frequently labeled by C1q in R+;AD+ brain. This correlated with less synapse loss and decreased phagocytosis of PSD95-labeled synaptic membranes by microglia. Together, these findings suggest that decreased Aβ plaque burden, as well as decreased neuron degeneration and synapse loss in R+;AD+ mice, contributed to the functional improvement in learning and memory behavior.

GWAS studies have identified >40 LOAD risk-associated polymorphic gene loci, which influence many genes expressed in microglia (64). While the impact of these polymorphisms on microglia function is only partially understood, two of the most important risk-associated variants APOE4 and TREM2^R47H^ appear to impair microglial immunometabolic function. For example, expression of APOE4 in microglia/CAMs reduces their accumulation around Aβ plaques, reduces expression of complement and lysosomal pathways in microglia, accelerates amyloid plaque burden and impairs synaptic plasticity (65). Similarly, TREM2 which is a high affinity receptor for Aβ and APOE4 (47, 66), is essential for transducing DAM activation response to Aβ (44). Microglia expressing the LOAD risk-associated variant TREM2^R47H^ have reduced affinity for their Aβ and APOE4 ligands, decreased Aβ plaque accumulation, and they poorly acquire a DAM activation state (67, 68). These phenotypes are associated with altered signaling from the TREM2/DAP12 complex that phosphorylates/activates the tyrosine kinase SYK to engage downstream signaling pathways mediated in part by mTOR, PI3K/AKT and mitochondrial oxidative phosphorylation (30, 32). Thus, emerging evidence points to SYK as an intracellular signaling hub downstream of TREM2 that drives microglial immunometabolic fitness and a neuroprotective phenotype in response to Aβ. Interestingly, we found clear evidence of increased SYK phosphorylation in ACE-expressing microglia in R+;AD+ PAM in response to Aβ, and that inhibition of SYK using a pharmacologic inhibitor or gene silencing abrogated the enhanced phagocytosis and endolysosomal trafficking of Aβ within ACE-expressing (R+;AD-) microglia. ACE-enhanced SYK signaling was also identified from snRNAseq pathway analysis that showed upregulation of several signaling pathways downstream of SYK in DAM, including mTOR, PI3/AKT and oxidative phosphorylation. Similarly, DSP pathway analysis in PAM from R+;AD+ mice identified increased translation and rRNA synthesis, known to be downstream of mTOR signaling, and increased c-lectin/CLEC7A receptor signaling, known to transduce Aβ binding and drive SYK phosphorylation and signaling (30). Taken together, these results indicate that ACE-expression in microglia increases SYK signaling and downstream pathway activation to enhance their AD protective function, and accordingly, the phenotypic effects of ACE expression in microglia are opposite to those observed in SYK-deficient microglia in 5xFAD mice (30, 32).

Microglia encircle Aβ plaques and they engage a DAM transcriptional activation state that regulates their immune response to process Aβ (44). In these studies, we identified two DAM populations in AD+ mice, DAM-A which appeared to represent classic DAM that consisted of both DAM stage I and DAM stage II microglia (44), and a much smaller DAM-B population. ACE expression in microglia eliminated nearly 50% of the DAM-B and increased the frequency of DAM-A, and it induced many genes in DAM-A related to enhanced endolysosomal trafficking, migration, inflammation, oxidative phosphorylation and SYK signaling. Microglia that engage the DAM transcriptional state are thought to represent PAM (44), although single cell/nuclei transcriptomic studies have not definitively resolved the spatial relationship of DAM with Aβ plaques. By using snRNAseq combined with DSP, we found a significant correlation between PAM gene signatures and those in DAM-A indicating they were associated with Aβ plaques, while DAM-B were apparently not plaque-associated. DAM-B appear to represent an Nkain2, Pde4b, St18 high expressing population of microglia that were previously described when APOE was ablated in neurons in a tauopathy mouse model (69). Future studies should further characterize the functional relevance of DAM-B in 5xFAD mice.

ACE is normally expressed in human and mouse neurons (70), but not in microglia (Fig. S1), raising an interesting question about whether ACE has any endogenous role in regulating microglia function in the brain. ACE is a membrane bound protein with two catalytic domains located outside of the cell membrane. Similar to cell surface APP, ACE is released from the cell surface by cleavage-secretion to produce soluble, catalytically active ACE under normal physiological conditions (71). While its function is very poorly understood in neurons, expression on the surface of neurons and cleavage-secretion could provide a trans-signaling mechanism to modulate microglia function. Hints that such a signaling mechanism may exist come from studies identifying altered microglial function in ACE haploinsufficient mice where neuronal ACE levels are decreased by 50% (26) and studies showing that ACE levels are increased in the brain from AD patients and in neurons treated with Aβ_42_ (70, 72). In the RACE mice developed for these studies, where ACE is abundantly expressed on the cell surface of microglia, it seems plausible that cleavage-secretion could enhance an ACE signaling pathway through local cis-signaling in microglia which may have evolved to respond to neuron-derived ACE cleavage products. Future studies should interrogate whether a neuron-microglia-ACE signaling pathway exists and if so, to identify the ACE cleavage product(s) that modulate the neuron protective mechanisms mediated by microglia. Presuming that future studies confirm a similar function of ACE in human microglia, ACE expression in human microglia may be useful to override immunometabolic defects caused by single or multiple LOAD risk-associated genes. Given that ACE-expressing microglia seem to complement signaling pathways that are impaired in microglia that express the major AD risk-associated proteins, such as TREM2^R47H^ and APOE4, it may be feasible to use ACE-expressing microglia in a cell-based therapy for AD. Promising cell-based therapies are in development for many diseases, and our recent first-in-human clinical trial demonstrated the safety profile of transplanted mesenchymal stem-cells expressing glial derived neurotrophic factor (GDNF) in the spinal cord for the treatment of ALS (73). Indeed, many pre-clinical studies are underway to engraft human microglia into mice as proof-of-concept that they may eventually be effective in human microglial transplantation therapy (74).

## METHODS

### Study Approval

All animal procedures were reviewed and approved by the Institutional Animal Care and Use Committee (IACUC) at Cedars Sinai Medical Center in Los Angeles California (IACUC approval: IACUC007292).

### Experimental Models

#### Mice and genotyping

##### Rosa-hACE-flox mice

Rosa-hACE-flox mice were generated using homologous recombination in C57BL/6 embryonic stem cells (ESC; MUBES-01001, Cyagen, Santa Clara, CA) to generate 100% ESC-derived founder mice using a targeting construct containing a self-deleting Neomycin phosphotransferase (sdNeo) positive-selection cassette (TurboKnockout, Cyagen, Santa Clara, CA). To generate the gene targeting construct, homologous recombination domains (HRDs) were generated using high fidelity Taq DNA polymerase (Q5 high-fidelity DNA polymerase, NEB, Ipswich, MA) and polymerase chain reaction (PCR) with a BAC plasmid containing the Gt(Rosa)26Sor gene sequence (RP23-401D9) as PCR DNA template. A 2647 bp 5’ HRD and a 4107 bp 3’ HRD flanking the XbaI site in intron 1 of the Rosa genomic locus was generated and the sdNeo-CAG-LSL-TurboGFP-P2A-hACE-3xF construct was cloned in reverse orientation into the Xba1 site flanked by the HRDs using isothermal assembly. The final targeting construct was verified using restriction nuclease mapping and Sanger sequencing. The targeting construct was linearized using AscI restriction endonuclease, electroporated into MUBES ESCs and selected for 24 hours after transfection in G418 (200 mg/mL). 187 clones were isolated and screened, and 27 candidate clones were identified using long range (LR) PCR across the 3’ HRD arm using primers PR1 and PR2. Additional PCR screening was performed to verify the presence of the Neo selection cassette and the SV40 poly adenylation sequence in the candidate clone ESC genome.

Southern blot analysis was performed on MfeI restricted genomic DNA from PCR confirmed candidate clones to verify appropriate genome targeting using a probe upstream of the 5’ HRD. The double stranded DNA probe was generated by PCR using BAC plasmid DNA (RP23-401D9) to amplify a ∼500 bp amplicon using primers PR3 and PR4. The probe DNA was labeled with Digoxigenin using DIG-High Prime DNA labeling kit and Southern blot detection was performed using DIG Nucleic Acid Detection kit and accompanying protocols. The restriction fragments hybridized to the probe were expected to generate a 12.6kb fragment on the wild type allele and a 10.3kb fragment on the targeted allele in MfeI restricted genomic DNA from heterozygous clones. Five ESC candidate clones identified by PCR (2A3, 2A4, 1B1, 1E2 and 2H3) were confirmed by Southern blotting analysis. Clone 2A3 was selected for blastocyst injection to generate 3 male and 2 female germline Rosa^+/hACE-flox^ mice. Founder mice were mated to C57BL/6J mice and heterozygotes were maintained on an isogenic C57BL/6J genetic background.

Rosa^+/hACE-flox^ mice were genotyped using genomic DNA isolated from tail biopsy or ear punch tissue with the following primers: OT2138 and OT2139 to generate a 247 bp Rosa wild type allele amplicon and OT2138 and OT2141 to generate a 451 bp hACE-flox knockin allele amplicon.

##### Cx3Cr1-CreERT2 mice

Heterozygous (Cx3Cr1^+/CreERT2^) mice were obtained from The Jackson Laboratory (Jax strain: #020940) and backcrossed to an isogenic C57BL/6J line. The mice were previously characterized to drive cre-recombinase expression in peripheral myeloid cells, microglia and CAMs. Approximately 21 days after intraperitoneal Tamoxifen injection peripheral myeloid cells are regenerated in their transgene-OFF state, but in microglia, which are permanently engrafted into the brain during embryogenesis, loxP site recombination endures in a transgene-ON state (38).

Mice were genotyped using genomic DNA isolated from tail biopsy tissue with the following primers: OT2011 and OT2012: to generate a 291 bp wild type allele amplicon and OT2011 and OT2013 to generate a 455 bp CreERT2 knockin allele amplicon.

##### 5xFAD mice

The original 5xFAD transgenic line, Tg6799 was obtained from The Jackson Laboratory (5xFAD^+/Tg6799^) and heterozygous mice were used in these studies. 5xFAD mice were previously generated to express five human familial amyloid mutations and they rapidly produce extracellular amyloid-beta (Aβ) deposition in the brain. 5xFAD mice exhibit cognitive and neurodegenerative changes, similar to Alzheimer’s Disease (39).

5xFAD mice were genotyped by using the previously identified transgene integration site to genotype heterozygous and homozygous mice by PCR (75). Mice were genotyped using genomic DNA isolated from tail biopsy or ear punch tissues with the following primers: OT2275 and OT2276 to generate a 402 bp wild type allele amplicon and OT2275 and OT2274 to generate a 274 bp transgenic allele amplicon.

##### RACE mice

RACE mice were generated by mating Cx3Cr1-CreERT2 mice to Rosa-hACE-flox mice. Mice with genotype Cx3Cr1^+/CreERT2^; Rosa^+/+^ were designated <u>RACE-</u> and mice with genotype Cx3Cr1^+/CreERT2^; Rosa^+/hACE-flox^ were designated <u>RACE+</u>.

All mouse lines were maintained on C57BL/6J isogenic background. Specific PCR reaction conditions for genotyping are available upon request.

## SUPPLEMENTAL INFORMATION

Supplemental methods and extended data can be found online.

## AUTHOR CONTRIBUTIONS

W.G.T. conceived the project and designed the studies. A.G., S.W., H.B., A.K.M., L.L and A.M. performed experiments, W.G.T, A.G., S.W., H.B., A.K.M., K.E.B and L.L. analyzed the data, and W.G.T. and A.G. wrote the manuscript.

## Supporting information

graphical abstract + supplemental content

## ACKNOWLEDGEMENTS

Supported by NIH: RF1-AG074365 (W.G.T.), R01-AI164519 (K.E.B), Cedars-Sinai Goldrich Alzheimer’s Grant (W.G.T, K.E.B). The authors thank Lin Karman for technical assistance and Dr. Alexandra Moser for reviewing the manuscript.

## REFERENCES

1. Association As. 2023: Alzheimer’s Disease Facts and Figures. 2023.

2. Frisoni GB, Altomare D, Thal DR, Ribaldi F, van der Kant R, Ossenkoppele R, et al. The probabilistic model of Alzheimer disease: the amyloid hypothesis revised. Nature reviews Neuroscience. 2022;23(1):53–66.

3. Clayton KA, Van Enoo AA, and Ikezu T. Alzheimer’s Disease: The Role of Microglia in Brain Homeostasis and Proteopathy. Front Neurosci. 2017;11:680.

4. Xu Y, Jin M-Z, Yang Z-Y, and Jin W-L. Microglia in neurodegenerative diseases. Neural Regeneration Research. 2021;16(2):270–80.

5. Chávez-Gutiérrez L, and Szaruga M. Mechanisms of neurodegeneration - Insights from familial Alzheimer’s disease. Semin Cell Dev Biol. 2020;105:75–85.

6. Wightman DP, Jansen IE, Savage JE, Shadrin AA, Bahrami S, Holland D, et al. A genome-wide association study with 1,126,563 individuals identifies new risk loci for Alzheimer’s disease. Nat Genet. 2021;53(9):1276–82.

7. Bellenguez C, Küçükali F, Jansen IE, Kleineidam L, Moreno-Grau S, Amin N, et al. New insights into the genetic etiology of Alzheimer’s disease and related dementias. Nature Genetics. 2022;54(4):412–36.

8. Kunkle BW, Grenier-Boley B, Sims R, Bis JC, Damotte V, Naj AC, et al. Genetic meta-analysis of diagnosed Alzheimer’s disease identifies new risk loci and implicates Abeta, tau, immunity and lipid processing. Nat Genet. 2019;51(3):414–30.

9. Miao J, Ma H, Yang Y, Liao Y, Lin C, Zheng J, et al. Microglia in Alzheimer’s disease: pathogenesis, mechanisms, and therapeutic potentials. Front Aging Neurosci. 2023;15:1201982.

10. Wingo AP, Liu Y, Gerasimov ES, Gockley J, Logsdon BA, Duong DM, et al. Integrating human brain proteomes with genome-wide association data implicates new proteins in Alzheimer’s disease pathogenesis. Nature Genetics. 2021;53(2):143–6.

11. Kehoe PG, Russ C, McIlory S, Williams H, Holmans P, Holmes C, et al. Variation in DCP1, encoding ACE, is associated with susceptibility to Alzheimer disease. Nat Genet. 1999;21(1):71–2.

12. Lehmann DJ, Cortina-Borja M, Warden DR, Smith AD, Sleegers K, Prince JA, et al. Large meta-analysis establishes the ACE insertion-deletion polymorphism as a marker of Alzheimer’s disease. American journal of epidemiology. 2005;162(4):305–17.

13. Marioni RE, Harris SE, Zhang Q, McRae AF, Hagenaars SP, Hill WD, et al. GWAS on family history of Alzheimer’s disease. Transl Psychiatry. 2018;8(1):99.

14. Meng Y, Baldwin CT, Bowirrat A, Waraska K, Inzelberg R, Friedland RP, et al. Association of polymorphisms in the Angiotensin-converting enzyme gene with Alzheimer disease in an Israeli Arab community. American journal of human genetics. 2006;78(5):871–7.

15. Esther CR, Marino EM, Howard TE, Machaud A, Corvol P, Capecchi MR, et al. The critical role of tissue angiotensin-converting enzyme as revealed by gene targeting in mice. J Clin Invest. 1997;99(10):2375–85.

16. Murray MD, Hendrie HC, Lane KA, Zheng M, Ambuehl R, Li S, et al. Antihypertensive Medication and Dementia Risk in Older Adult African Americans with Hypertension: A Prospective Cohort Study. Journal of General Internal Medicine. 2018;33(4):455–62.

17. Hughes D, Judge C, Murphy R, Loughlin E, Costello M, Whiteley W, et al. Association of Blood Pressure Lowering With Incident Dementia or Cognitive Impairment: A Systematic Review and Meta-analysis. JAMA : the journal of the American Medical Association. 2020;323(19):1934–44.

18. Ding J, Davis-Plourde KL, Sedaghat S, Tully PJ, Wang W, Phillips C, et al. Antihypertensive medications and risk for incident dementia and Alzheimer’s disease: a meta-analysis of individual participant data from prospective cohort studies. The Lancet Neurology. 2020;19(1):61–70.

19. Li N-C, Lee A, Whitmer RA, Kivipelto M, Lawler E, Kazis LE, et al. Use of angiotensin receptor blockers and risk of dementia in a predominantly male population: prospective cohort analysis. BMJ (Clinical research ed). 2010;340:b5465.

20. van Dalen JW, Marcum ZA, Gray SL, Barthold D, Moll van Charante EP, van Gool WA, et al. Association of Angiotensin II–Stimulating Antihypertensive Use and Dementia Risk. Neurology. 2021;96(1):e67–e80.

21. Barthold D, Joyce G, Wharton W, Kehoe P, and Zissimopoulos J. The association of multiple anti-hypertensive medication classes with Alzheimer’s disease incidence across sex, race, and ethnicity. PLOS ONE. 2018;13(11):e0206705.

22. Rao A, Bhat SA, Shibata T, Giani JF, Rader F, Bernstein KE, et al. Diverse biological functions of the renin-angiotensin system. Med Res Rev. 2024;44(2):587–605.

23. Sinka L, Biasch K, Khazaal I, Peault B, and Tavian M. Angiotensin-converting enzyme (CD143) specifies emerging lympho-hematopoietic progenitors in the human embryo. Blood. 2012;119(16):3712–23.

24. Bernstein KE, Khan Z, Giani JF, Cao DY, Bernstein EA, and Shen XZ. Angiotensin-converting enzyme in innate and adaptive immunity. Nature reviews Nephrology. 2018;14(5):325–36.

25. Zou K, Yamaguchi H, Akatsu H, Sakamoto T, Ko M, Mizoguchi K, et al. Angiotensin-Converting Enzyme Converts Amyloid-beta protein Ab 1-42 to Ab 1-40, and Its Inhibition Enhances Brain Ab Deposition. Journal of Neuroscience. 2007;27(32):8628–35.

26. Liu S, Ando F, Fujita Y, Liu J, Maeda T, Shen X, et al. A clinical dose of angiotensin-converting enzyme (ACE) inhibitor and heterozygous ACE deletion exacerbate Alzheimer’s disease pathology in mice. J Biol Chem. 2019;294(25):9760–70.

27. Shen XZ, Li P, Weiss D, Fuchs S, Xiao HD, Adams JA, et al. Mice with enhanced macrophage angiotensin-converting enzyme are resistant to melanoma. Am J Pathol. 2007;170(6):2122–34.

28. Khan Z, Shen XZ, Bernstein EA, Giani JF, Eriguchi M, Zhao TV, et al. Angiotensin-converting enzyme enhances the oxidative response and bactericidal activity of neutrophils. Blood. 2017;130(3):328–39.

29. Okwan-Duodu D, Weiss D, Peng Z, Veiras LC, Cao DY, Saito S, et al. Overexpression of myeloid angiotensin-converting enzyme (ACE) reduces atherosclerosis. Biochemical and biophysical research communications. 2019;520(3):573–9.

30. Wang S, Sudan R, Peng V, Zhou Y, Du S, Yuede CM, et al. TREM2 drives microglia response to amyloid-β via SYK-dependent and -independent pathways. Cell. 2022;185(22):4153–69.e19.

31. Schafer DP, and Stillman JM. Microglia are SYK of Aβ and cell debris. Cell. 2022;185(22):4043–5.

32. Ennerfelt H, Frost EL, Shapiro DA, Holliday C, Zengeler KE, Voithofer G, et al. SYK coordinates neuroprotective microglial responses in neurodegenerative disease. Cell. 2022;185(22):4135–52.e22.

33. Bernstein KE, Gonzalez-Villalobos RA, Giani JF, Shah K, Bernstein E, Janjulia T, et al. Angiotensin-converting enzyme overexpression in myelocytes enhances the immune response. Biol Chem. 2014;395(10):1173–8.

34. Soriano P. Generalized lacZ expression with the ROSA26 Cre reporter strain. Nat Genet. 1999;21(1):70–1.

35. Srinivas S, Watanabe T, Lin CS, William CM, Tanabe Y, Jessell TM, et al. Cre reporter strains produced by targeted insertion of EYFP and ECFP into the ROSA26 locus. BMC developmental biology. 2001;1(1):4.

36. Yona S, Kim KW, Wolf Y, Mildner A, Varol D, Breker M, et al. Fate mapping reveals origins and dynamics of monocytes and tissue macrophages under homeostasis. Immunity. 2013;38(1):79–91.

37. McKinsey GL, Lizama CO, Keown-Lang AE, Niu A, Santander N, Larpthaveesarp A, et al. A new genetic strategy for targeting microglia in development and disease. eLife. 2020;9:e54590.

38. Parkhurst CN, Yang G, Ninan I, Savas JN, Yates JR, 3rd, Lafaille JJ, et al. Microglia promote learning-dependent synapse formation through brain-derived neurotrophic factor. Cell. 2013;155(7):1596–609.

39. Oakley H, Cole SL, Logan S, Maus E, Shao P, Craft J, et al. Intraneuronal beta-amyloid aggregates, neurodegeneration, and neuron loss in transgenic mice with five familial Alzheimer’s disease mutations: potential factors in amyloid plaque formation. J Neurosci. 2006;26(40):10129–40.

40. Ohno M. Failures to reconsolidate memory in a mouse model of Alzheimer’s disease. Neurobiol Learn Mem. 2009;92(3):455–9.

41. Ferro A, Auguste YSS, and Cheadle L. Microglia, Cytokines, and Neural Activity: Unexpected Interactions in Brain Development and Function. Front Immunol. 2021;12:703527.

42. Stence N, Waite M, and Dailey ME. Dynamics of microglial activation: a confocal time-lapse analysis in hippocampal slices. Glia. 2001;33(3):256–66.

43. Navarro V, Sanchez-Mejias E, Jimenez S, Munoz-Castro C, Sanchez-Varo R, Davila JC, et al. Microglia in Alzheimer’s Disease: Activated, Dysfunctional or Degenerative. Front Aging Neurosci. 2018;10:140.

44. Keren-Shaul H, Spinrad A, Weiner A, Matcovitch-Natan O, Dvir-Szternfeld R, Ulland TK, et al. A Unique Microglia Type Associated with Restricting Development of Alzheimer’s Disease. Cell. 2017;169(7):1276–90 e17.

45. Herre J, Marshall ASJ, Caron E, Edwards AD, Williams DL, Schweighoffer E, et al. Dectin-1 uses novel mechanisms for yeast phagocytosis in macrophages. Blood. 2004;104(13):4038–45.

46. Krasemann S, Madore C, Cialic R, Baufeld C, Calcagno N, El Fatimy R, et al. The TREM2-APOE Pathway Drives the Transcriptional Phenotype of Dysfunctional Microglia in Neurodegenerative Diseases. Immunity. 2017;47(3):566–81.e9.

47. Zhao Y, Wu X, Li X, Jiang LL, Gui X, Liu Y, et al. TREM2 Is a Receptor for beta-Amyloid that Mediates Microglial Function. Neuron. 2018;97(5):1023–31.e7.

48. Roy ER, Chiu G, Li S, Propson NE, Kanchi R, Wang B, et al. Concerted type I interferon signaling in microglia and neural cells promotes memory impairment associated with amyloid beta plaques. Immunity. 2022;55(5):879–94 e6.

49. Eimer WA, and Vassar R. Neuron loss in the 5XFAD mouse model of Alzheimer’s disease correlates with intraneuronal Abeta42 accumulation and Caspase-3 activation. Molecular neurodegeneration. 2013;8:2.

50. Hong S, Beja-Glasser VF, Nfonoyim BM, Frouin A, Li S, Ramakrishnan S, et al. Complement and microglia mediate early synapse loss in Alzheimer mouse models. Science. 2016;352(6286):712–6.

51. Roy ER, Wang B, Wan Y-W, Chiu GS, Cole AL, Yin Z, et al. Type I interferon response drives neuroinflammation and synapse loss in Alzheimer disease. The Journal of Clinical Investigation. 2020.

52. Huang Y, Happonen KE, Burrola PG, O’Connor C, Hah N, Huang L, et al. Microglia use TAM receptors to detect and engulf amyloid β plaques. Nature immunology. 2021;22(5):586–94.

53. Chen X, and Holtzman DM. Emerging roles of innate and adaptive immunity in Alzheimer’s disease. Immunity. 2022;55(12):2236–54.

54. Palmieri M, Impey S, Kang H, di Ronza A, Pelz C, Sardiello M, et al. Characterization of the CLEAR network reveals an integrated control of cellular clearance pathways. Human Molecular Genetics. 2011;20(19):3852–66.

55. Settembre C, and Medina DL. TFEB and the CLEAR network. Methods Cell Biol. 2015;126:45–62.

56. Torres-Hernandez A, Wang W, Nikiforov Y, Tejada K, Torres L, Kalabin A, et al. Targeting SYK signaling in myeloid cells protects against liver fibrosis and hepatocarcinogenesis. Oncogene. 2019;38(23):4512–26.

57. Cao DY, Spivia WR, Veiras LC, Khan Z, Peng Z, Jones AE, et al. ACE overexpression in myeloid cells increases oxidative metabolism and cellular ATP. J Biol Chem. 2020;295(5):1369–84.

58. Schweig JE, Yao H, Beaulieu-Abdelahad D, Ait-Ghezala G, Mouzon B, Crawford F, et al. Alzheimer’s disease pathological lesions activate the spleen tyrosine kinase. Acta Neuropathol Commun. 2017;5(1):69.

59. Makhoul S, Dorschel S, Gambaryan S, Walter U, and Jurk K. Feedback Regulation of Syk by Protein Kinase C in Human Platelets. International journal of molecular sciences. 2019;21(1).

60. Tsang E, Giannetti AM, Shaw D, Dinh M, Tse JK, Gandhi S, et al. Molecular mechanism of the Syk activation switch. J Biol Chem. 2008;283(47):32650–9.

61. Ryan DK, Karhunen V, Su B, Traylor M, Richardson TG, Burgess S, et al. Genetic Evidence for Protective Effects of Angiotensin-Converting Enzyme Against Alzheimer Disease But Not Other Neurodegenerative Diseases in European Populations. Neurology Genetics. 2022;8(5):e200014.

62. Okwan-Duodu D, Datta V, Shen XZ, Goodridge HS, Bernstein EA, Fuchs S, et al. Angiotensin-converting enzyme overexpression in mouse myelomonocytic cells augments resistance to Listeria and methicillin-resistant Staphylococcus aureus. J Biol Chem. 2010;285(50):39051–60.

63. Bernstein KE, Koronyo Y, Salumbides BC, Sheyn J, Pelissier L, Lopes DH, et al. Angiotensin-converting enzyme overexpression in myelomonocytes prevents Alzheimer’s-like cognitive decline. J Clin Invest. 2014;124(3):1000–12.

64. Sudwarts A, and Thinakaran G. Alzheimer’s genes in microglia: a risk worth investigating. Molecular neurodegeneration. 2023;18(1):90.

65. Liu CC, Wang N, Chen Y, Inoue Y, Shue F, Ren Y, et al. Cell-autonomous effects of APOE4 in restricting microglial response in brain homeostasis and Alzheimer’s disease. Nature immunology. 2023;24(11):1854–66.

66. Atagi Y, Liu CC, Painter MM, Chen XF, Verbeeck C, Zheng H, et al. Apolipoprotein E Is a Ligand for Triggering Receptor Expressed on Myeloid Cells 2 (TREM2). J Biol Chem. 2015;290(43):26043–50.

67. Song WM, Joshita S, Zhou Y, Ulland TK, Gilfillan S, and Colonna M. Humanized TREM2 mice reveal microglia-intrinsic and -extrinsic effects of R47H polymorphism. J Exp Med. 2018;215(3):745–60.

68. Zhou Y, Song WM, Andhey PS, Swain A, Levy T, Miller KR, et al. Human and mouse single-nucleus transcriptomics reveal TREM2-dependent and TREM2-independent cellular responses in Alzheimer’s disease. Nature medicine. 2020;26(1):131–42.

69. Koutsodendris N, Blumenfeld J, Agrawal A, Traglia M, Grone B, Zilberter M, et al. Neuronal APOE4 removal protects against tau-mediated gliosis, neurodegeneration and myelin deficits. Nature Aging. 2023;3(3):275–96.

70. Cuddy LK, Prokopenko D, Cunningham EP, Brimberry R, Song P, Kirchner R, et al. Abeta-accelerated neurodegeneration caused by Alzheimer’s-associated ACE variant R1279Q is rescued by angiotensin system inhibition in mice. Sci Transl Med. 2020;12(563).

71. Ramchandran R, Sen GC, Misono K, and Sen I. Regulated cleavage-secretion of the membrane-bound angiotensin-converting enzyme. J Biol Chem. 1994;269(3):2125–30.

72. Miners S, Ashby E, Baig S, Harrison R, Tayler H, Speedy E, et al. Angiotensin- converting enzyme levels and activity in Alzheimer’s disease: differences in brain and CSF ACE and association with ACE1 genotypes. Am J Transl Res. 2009;1(2):163–77.

73. Baloh RH, Johnson JP, Avalos P, Allred P, Svendsen S, Gowing G, et al. Transplantation of human neural progenitor cells secreting GDNF into the spinal cord of patients with ALS: a phase 1/2a trial. Nature medicine. 2022;28(9):1813–22.

74. Ifediora N, Canoll P, and Hargus G. Human stem cell transplantation models of Alzheimer’s disease. Front Aging Neurosci. 2024;16:1354164.

75. Goodwin LO, Splinter E, Davis TL, Urban R, He H, Braun RE, et al. Large-scale discovery of mouse transgenic integration sites reveals frequent structural variation and insertional mutagenesis. Genome research. 2019;29(3):494–505.

