## Supplementary material for "Angiotensin Converting Enzyme (ACE) expression in microglia reduces amyloid β deposition and neurodegeneration by increasing SYK signaling and endolysosomal trafficking": graphical abstract + supplemental content

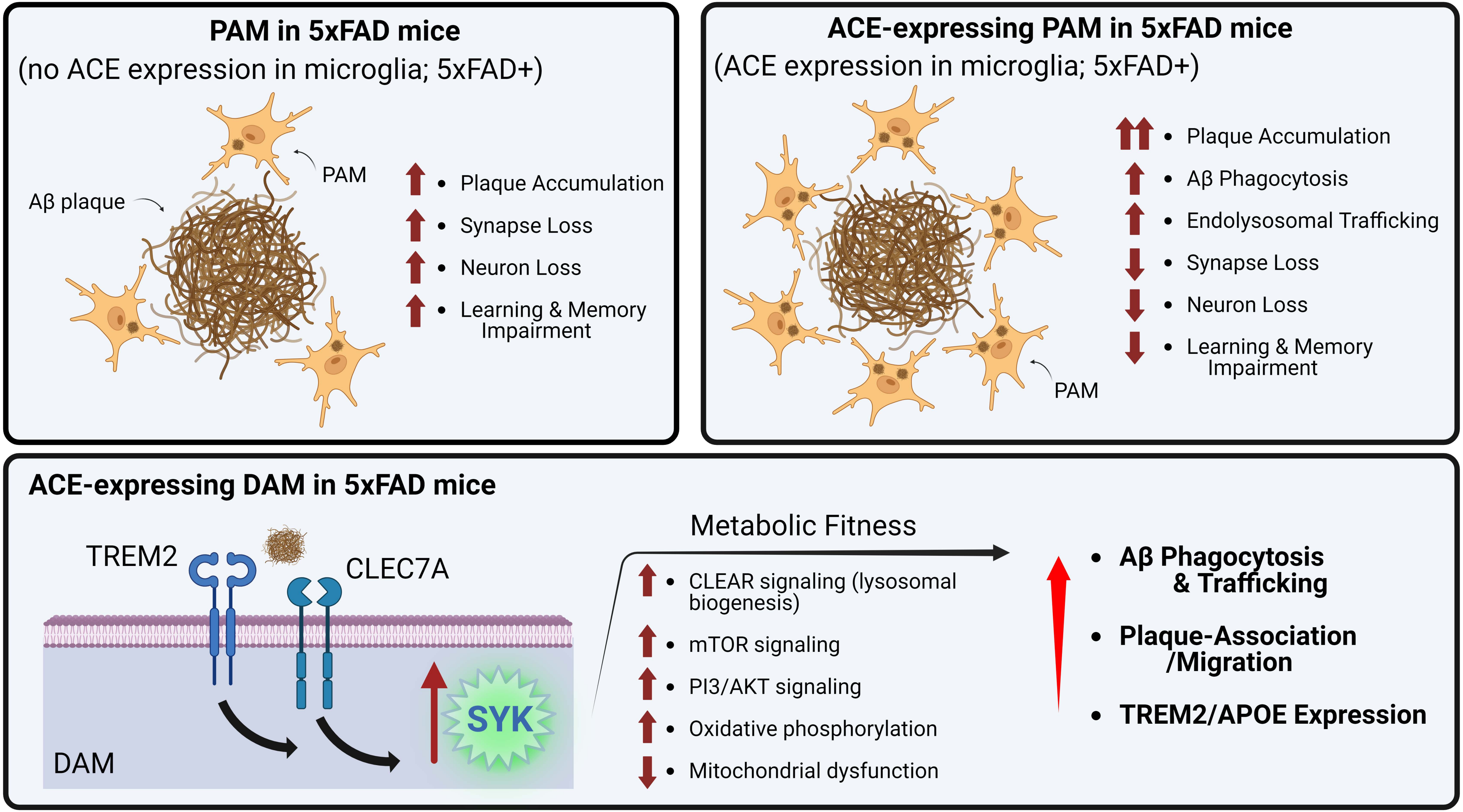

PAM in 5xFAD mice
(no ACE expression in microglia; 5xFAD+)
Aβ plaque
PAM
Plaque Accumulation
Synapse Loss
Neuron Loss
Learning & Memory Impairment
ACE-expressing PAM in 5xFAD mice
(ACE expression in microglia; 5xFAD+)
Plaque Accumulation
Aβ Phagocytosis
Endolysosomal Trafficking
Synapse Loss
Neuron Loss
Learning & Memory Impairment
PAM
ACE-expressing DAM in 5xFAD mice
TREM2
CLEC7A
SYK
DAM
Metabolic Fitness
CLEAR signaling (lysosomal biogenesis)
mTOR signaling
PI3/AKT signaling
Oxidative phosphorylation
Mitochondrial dysfunction
Aβ Phagocytosis & Trafficking
Plaque-Association /Migration
TREM2/APOE Expression

### Microglial angiotensin converting enzyme (ACE) reduces pathologic amyloid β deposition and neurodegeneration by increasing SYK signaling and endolysosomal trafficking.

Andrew R. Gomez, Hyae Ran Byun, Shaogen Wu, AKM Ghulam Mohammad, Jasmine Ikbariyeh, Jaelin Chen, Alek Muro, Lin Li, Ken E. Bernstein, Richard Ainsworth and Warren G. Tourtellotte

#### In Brief

Expression of angiotensin converting enzyme (ACE) in microglia potentiates their immune function by increasing expression of TREM2 and enhancing SYK signaling. ACE expression in microglia in 5xFAD mice enhanced their migration to Aβ plaques, and their phagocytosis and endolysosomal trafficking of Aβ which results in decreased amyloid plaque burden, decreased neuron and synapse loss, and rescue of learning and memory behavioral abnormalities in the AD model mice.

#### Highlights

- Expression of ACE in microglia leads to decreased amyloid plaque burden, neuron and synapse loss and rescue of learning and memory abnormalities in 5xFAD model mice.
- ACE expression potentiates SYK signaling to enhance microglial metabolic fitness by potentiating inducing TREM2 expression, lysosomal biogenesis pathways (CLEAR signaling), mTOR and PI3/AKT signaling, and oxidative phosphorylation.
- ACE expressing microglia could be a candidate for cell-based microglia replacement therapy in humans with AD.

### Supplemental Methods and Extended Data

Angiotensin Converting Enzyme (ACE) expression in microglia reduces amyloid β deposition and neurodegeneration by increasing SYK signaling and endolysosomal trafficking.

Andrew R. Gomez, Hyae Ran Byun, Shaogen Wu, AKM Ghulam Muhammad, Jasmine Ikbariyeh, Jaelin Chen, Alek Muro, Lin Li, Kenneth E. Bernstein, Richard Ainsworth and Warren G. Tourtellotte

#### Supplemental Methods.

##### Key Resources Table

| REAGENT or RESOURCE | SOURCE | IDENTIFIER |
|---|---|---|
| **Antibodies and Labeling reagents** | | |
| Normal Donkey Serum | Equitech | DS05 |
| Fc Receptor Blocker (1:50) | Biolegend | 101319 |
| Thioflavin S | Sigma-Aldrich | 76264-656 |
| Amylo-Glo RTD | VWR | TR-300-AG |
| DAPI DNA staining solution | Abcam | ab228549 |
| SYTO Green nucleic acid stain | ThermoFisher Scientific | S7575 |
| P505 SYK inhibitor (PRT062607) | MedChemExpress | HY-15323 |
| PRIMARY ANTIBODIES | | |
| Mouse anti-Cx3Cr1, PE conj. (1:10) | R&D systems | FAB5825P |
| Mouse anti-PSD95 (1:100) | Millipore-Sigma | MABN68 |
| Rat anti-CD11b, BV605 conj. (1:100) | Biolegend | 101237 |
| Rat anti-mouse, Total-seq, Hashtag B-0301 | Biolegend | 155831 |
| Rat anti-mouse, Total-seq, Hashtag B-0302 | Biolegend | 155833 |
| Rat anti-mouse, Total-seq, Hashtag B-0303 | Biolegend | 155835 |
| Rat anti-mouse, Total-seq, Hashtag B-0304 | Biolegend | 155837 |
| Rat anti-mouse, Total-seq, Hashtag B-0305 | Biolegend | 155839 |
| Rat anti-mouse, Total-seq, Hashtag B-0306 | Biolegend | 155841 |
| Rabbit anti-Iba1, Alexa 647 conj. (1:100) | Cell Signaling | 78060S |
| Mouse anti-Aβ, Alexa 488 conj. (1:100) | Biolegend | 803013 |
| Goat anti-Iba1 (1:500) | ThermoFisher Scientific | PA5-18039 |
| Goat anti-human ACE (1:500) | ThermoFisher Scientific | PA5-47964 |
| Rabbit anti-ACE (1:500) | Abcam | ab75762 |
| Rabbit anti-$A\beta_{1-42}$ (1:250) | Abcam | ab201060 |
| Rabbit anti-Iba1 (1:500) | Wako | 016-20001 |
| Rabbit anti-CD68 (1:500) | Abcam | ab125212 |
| Rabbit anti-C1q (1:1000) | Abcam | Ab182451 |
| Rabbit anti-pSyk, Y352 (1:200) | Cell Signaling | CST2717 |
| Rabbit anti-NeuN, Alexa 647 conj. (1:100) | Abcam | ab190565 |
| Rabbit anti-Sox10, Alexa 647 conj. (1:100) | Abcam | ab270151 |
| Rabbit anti-Sox9, Alexa 488 conj. (1:100) | Abcam | ab196450 |
| Guinea Pig anti-vGLUT1 (1:1000) | Millipore-Sigma | AB5905 |
| Guinea Pig anti NeuN (1:1000) | Millipore-Sigma | ABN90 |

| | | |
|---|---|---|
| SECONDARY ANTIBODIES | | |
| Donkey anti-mouse IgG (H+L), DyLight 649 | Jackson Immunoresearch | 715-495-150 |
| Donkey anti-mouse IgG (H+L), Alexa 488 | Thermofisher Scientific | A21202 |
| Donkey anti-mouse IgG (H+L), Alexa 647 | Thermofisher Scientific | A31571 |
| Donkey anti-rabbit IgG (H+L), Alexa 568 | Thermofisher Scientific | A10042 |
| Donkey anti-rabbit IgG (H+L), Alexa 647 | Thermofisher Scientific | A31573 |
| Donkey anti-rabbit IgG (H+L), Alexa 488 | Thermofisher Scientific | A21206 |
| Donkey anti-goat IgG (H+L), Alexa 647 | Thermofisher Scientific | A21447 |
| Donkey anti-goat IgG (H+L), Alexa 488 | Thermofisher Scientific | A32814 |
| Donkey anti-goat IgG (H+L), Alexa 568 | Thermofisher Scientific | A11057 |
| Donkey anti-guinea pig (H+L), Cy3 | Jackson Immunoresearch | 706-165-148 |
| Donkey anti-guinea pig (H+L), Alexa 647 | Jackson Immunoresearch | 706-605-148 |

| **Chemicals, peptides and recombinant proteins** | | |
|---|---|---|
| Tamoxifen | Sigma-Aldrich | T5648 |
| 4-hydroxy Tamoxifen (4-OHT) | Sigma-Aldrich | H7904 |
| A$\beta_{1\text{-}42}$ peptide | Anaspec | AS-20276 |
| A$\beta_{1\text{-}42}$, Hylite 647, conj. | Anaspec | AS-64161 |
| RNase Inhibitor | New England Biolabs | M0314S |
| DMEM, high glucose medium | ThermoFisher Scientific | 11965118 |
| HBSS medium | ThermoFisher Scientific | 14175095 |
| Recombinant murine GM-CSF | PeproTech | 315-03 |
| Recombinant murine M-CSF | PeproTech | 315-02 |
| G418 Sulfate (Geneticin) | ThermoFisher Scientific | 11811098 |
| 4’ ,6-Diamidino-2-phenylindole (DAPI) | Life Technologies | D21490 |
| MACS buffer | Miltenyi | 130-092-987 |
| RIPA buffer | ThermoFisher Scientific | 32955 |
| TRIzol | Invitrogen | 16096020 |
| BCA Protein Assay | ThermoFisher Scientific | 23228 |
| Triton X-100 | Sigma-Aldrich | X100 |
| SDS | Fisher Scientific | BP166 |
| EDTA | Fisher Scientific | S311 |
| DMSO | Sigma-Aldrich | D1435 |
| Acetone | Sigma-Aldrich | 1006801 |
| ProLong Gold, hard set mounting media | ThermoFisher Scientific | P36930 |
| Percoll Density Gradient Media | Fisher Scientific | 45001748 |
| DNase I | ThermoFisher Scientific | EN0521 |
| TLCK Trypsin Inhibitor | Santa Cruz Biotechnology | sc-201296 |
| HEPES | Thermofisher Scientific | 15630080 |
| Paraformaldehyde | Millipore-Sigma | P6148 |
| Ketamine (100 mg/mL) | MWI Animal Health | 501072 |
| Xylazine (20 mg/mL) | MWI Animal Health | 003437 |
| Protease and phosphatase inhibitor | Roche | 1836153 |
| pHrodo-Red Zymosan bioparticles | Thermofisher Scientific | P35364 |

| | | |
|---|---|---|
| SMARTpool (Mouse Syk) | Dharmacon Research | L-041084-00 |
| SMARTpool (non-targeting control/scramble) | Dharmacon Research | D-001810-10-05 |
| DharmaFECT transfection reagent | Dharmacon Research | T-2005-01 |
| **Plasmids, DNA primers and probes** | | |
| Murine BAC plasmid | BACPAC Genomics | RP23-401D9 |
| GACTAGAGCTTGCGGAACCCTT | IDT DNA | PR1 |
| GGAGCCATTCAGTGTTCACTATGT | IDT DNA | PR2 |
| GGAGAGGCGTTCAGGAAGATTATGG | IDT DNA | PR3 |
| CAGCTACCTTTACACACCATTGCACC | IDT DNA | PR4 |
| CCACATCAAGTTCTATCTAGGA | IDT DNA | OT2138 |
| TAGCAAACAAGAGACCACAT | IDT DNA | OT2139 |
| TTTTTTGTGTCTCTCACTCG | IDT DNA | OT2141 |
| CACGTGATCTGGTTTGCTGC | IDT DNA | OT2011 |
| CACCAGACCGAACGTGAA | IDT DNA | OT2012 |
| GACCGACGATGAAGCATG | IDT DNA | OT2013 |
| CGGGCCTCTTCGCTATTAC | IDT DNA | OT2274 |
| CTCTGTAAACACATCACAGCATAAA | IDT DNA | OT2275 |
| GACAGGAGTGAAGTCCCAAAG | IDT DNA | OT2276 |
| GAAGCCTTGCTAAGTGCGACA | IDT DNA | OT2449 |
| AAGTGCCGTGAATGGGTGAC | IDT DNA | OT2450 |
| CATCTTCCAGGAGCGAGACC | IDT DNA | OT2336 |
| GGCGGAGATGATGACCCTTT | IDT DNA | OT2337 |
| GeoMx, Mouse Whole Transcriptome Atlas | Nanostring | GMX-RNA-NGS-MsWTA |
| **Commercial Assays and kits** | | |
| DIG-High Prime DNA labeling kit | Millipore Sigma | 11745832910 |
| DIG-Nucleic Acid Detection kit | Roche | 11175041910 |
| pHrodo-Red, succinimidyl ester | Thermofisher Scientific | P36600 |
| Nuclei EZ Nuclei Isolation kit | Millipore-Sigma | NUC101-1KT |
| V-PLEX Mouse Cytokine Assay kit | Meso Scale Discovery | K15245D-1 |
| V-PLEX Aβ Peptide Panel (6E10) kit | Meso Scale Discovery | K15200E-1 |
| Chromium GEM Chip G | 10X Genomics | 1000127 |
| Chromium NEXT GEM 3’ Kit (v.3.1) | 10X Genomics | 1000269 |
| 3’ Feature Bar Code Kit | 10X Genomic | 1000262 |
| Dual Index Kit NT Set A | 10X Genomics | 1000242 |
| Superscript IV VILO Master Mix | ThermoFisher | 11766050 |
| PowerTrack SYBR Green Master Mix | ThermoFisher | A46110 |
| **Experimental models: Cell Lines** | | |
| Primary murine microglia | This paper | N/A |
| Murine ESC | Cyagan, Santa Clara, CA | MUBES-01001 |
| **Experimental models: Organisms/strains** | | |

| | | |
|---|---|---|
| Rosa-hACE-flox mice | This paper | N/A |
| RACE mice | This paper | N/A |
| Cx3Cr1-CreERT2 mice | Jackson Laboratories | 020940 |
| 5xFAD (Tg6799) mice | Jackson Laboratories | 008730 |
| C57BL/6J mice | Jackson Laboratories | 000664 |
| **Deposited Data** | | |
| GeoMx (Nanostring) | Nanostring, this paper | N/A |
| snRNAseq (10x Genomics) | 10x Genomics, this paper | N/A |
| **Software** | | |
| Imaris Analysis Software | Oxford Instruments | v. 10.1 |
| Zen Control and Image Analysis Software | Zeiss, Inc. | v. 2.3 |
| IncuCyte Control Software | Sartorius | v. 2022B, Rev3 |
| IncuCyte Analysis Software | Sartorius | v. 2022B, Rev2 |
| Graphpad Prism Statistical Analysis Software | Dotmatics | v. 10 |
| FlowJo Software | BD Bioscience | v. 10.10 |
| ANY-maze software | Stoelting Co. | v. 7.35 |
| GeoMx DSP control and analysis software | Nanostring | v. 2.3.0.268 |
| Reactome Pathway Database | Reactome | v. 78 |
| Ingenuity Pathway Analysis Software | Qiagen | v. Release 2023 |
| Cell Ranger Software | 10X Genomics | v. 6.1.2 |
| Image J | NIH | v. 1.54h |

#### Tissue Processing

Mice were euthanized with 80 mg/kg Ketamine/20 mg/kg Xylazine after IP injection, thoracotomy and transcardial perfusion with 20mL of Phosphate buffered saline (PBS; 100 mM, pH = 7.4). One brain hemisphere was post-fixed overnight in 4% paraformaldehyde in PBS (PFA), then transferred to PBS, 0.01% sodium azide for extended storage at 4 °C. The contralateral hemisphere was dissected to isolate hippocampal, cortical and thalamic structures, flash frozen, and stored at -80 °C. For protein extraction, unfixed frozen tissues were cryopulverized in RIPA buffer, lysed by passage through a 25G needle and centrifuged at 14,000 x *g*. The concentration of soluble protein from the supernatant was determined by BCA protein assay according to the manufacture's protocol.

#### Primary Microglia Isolation and Culture

Primary microglia were isolated from the cortex of neonatal (P1-P2) RACE- and RACE+ mice. Minced cortical tissues were dissociated by trituration and mixed astrocyte/microglia cultures were established in high glucose DMEM media containing of 10 ng/ml GM-CSF and 10 ng/ml M-CSF (GM) for 1 week. The cells were then treated with 2 μM 4-hydroxy Tamoxifen (4-OHT) for 3 days to induce ACE expression in RACE+, but not RACE- cells. To obtain primary microglia, the mixed culture was shaken at 220 rpm at 37 °C for 3 hours, the floating cells were recovered and centrifuged at 1000 x *g* for 3 min., the cell pellet was resuspended with GM and the cells were seeded onto culture plates or chamber slides. Primary microglia were maintained at 37 °C in high glucose DMEM under 5% CO2 until experiments were performed.

Pharmacological inhibition of SYK: Primary microglia were pretreated for 1 hour with 1 μM of the SYK tyrosine kinase inhibitor, P505 (PRT062607) prior to treatment with HiLyte-647 labeled

$oAb_{1-42}$ for 2 hours.

Syk mRNA silencing: Syk mRNA was targeted for silencing SMARTpool for Syk (L-041084-00) and control non-targeting pool/scramble (D-001810-10-05) from Dharmacon Research. Primary microglia were transfected with 25nM siRNA using DharmaFECT transfection reagent an then 48 hours later were treated with HiLyte-647 labeled $oAb_{1-42}$ for 2 hours.

**Total RNA extraction and qPCR**

Total RNA was isolated from primary microglia using TRIzol reagent (Invitrogen) according to manufacturer's specifications. Complementary DNA (cDNA) was synthesized using Superscript IV VILO Master Mix and the manufacturer's protocol for cDNA synthesis (ThermoFisher). qPCR was performed on a StepOne Plus real-time PCR instrument (ABI) using SYBR green master mix (Invitrogen). Mouse Syk with primers OT2449 and OT2450 and mouse GAPDH with primers OT2336 and OT2337 were used to amplify cDNA. Mouse GAPDH was used to normalize the samples for relative mRNA quantification using the $\Delta\Delta C_T$ method.

**Immunofluorescence**

Tissues: Coronal brain tissue sections (50 µm thickness) were generated using a vibratome (Leica VT1000S) and stored in PBS, 0.01% sodium azide at 4 °C. Prior to antibody treatment, free floating sections were washed in PBS for 10 min., cleared in an acetone-H2O series of solutions (25%-50%-25%) for 20 min. each, and permeabilized in PBS containing 10 mM EDTA, 0.2%Triton-X100, 0.1% SDS and 10%DMSO for 1 hr.

Cells: Prior to antibody treatment, cultured cells were fixed in PFA for 10 min., washed in PBS for 10 min., and permeabilized by treatment with PBS containing 0.2% Triton X-100 for 10 min.

Immunofluorescence was performed on floating sections or cultured cells by first blocking non-specific antibody binding using 5% normal donkey serum in PBS for 1 hour. Primary antibody incubation was performed overnight at room temperature (RT) or 4 °C. To label amyloid in plaques, Rabbit anti-$A\beta_{1-42}$, Thioflavin S or Amylo-Glo was used according to manufacturer's specifications. To label microglia, rabbit or goat anti-Iba1 antibodies were used. To label the intracellular endolysosomal compartment, rabbit anti-CD68 antibody was used. To label post-synaptic specializations, mouse anti-PSD95 was used and to label pre-synaptic excitatory axon terminals, a guinea pig anti-vGLUT1 antibody was used. To label compliment tagging in post-synaptic receptors, a rabbit anti-C1q antibody was used. To label neuronal nuclei, a guinea pig anti-NeuN antibody was used. To label activated Syk kinase, a rabbit anti-phospho-SYK (Y352) antibody was used.

After primary antibody incubation at RT or 4 °C overnight, tissue sections were washed in PBS and incubated for 2 hours with appropriate secondary antibodies (Donkey anti-mouse, anti-goat, anti-rabbit, anti-guinea pig) conjugated to fluorophores and diluted 1:1000 in PBS. Sections were washed in PBS and mounted on glass slides and coverslipped using hard setting mounting media. In some experiments, sections were stained with DAPI or SYTO Green to label nuclei according to the manufacturers protocol prior to coverslipping.

**Tamoxifen Treatment**

Mice: Tamoxifen (TMX) was prepared at 20 mg/mL and dissolved in corn oil. Intraperitoneal injections (125 mg/kg, IP) for RACE- and RACE+ mice were performed daily starting at postnatal day 28 for four consecutive days, each separated by a 24-hour interval. Tamoxifen treated RACE- and RACE+ mice were designated R- and R+ mice, respectively.

Cells: Primary microglia from RACE- and RACE+ mice were treated with 2µM 4-hydroxy-tamoxifen (4-OHT) for 3 days to recombine the LSL cassette in the genome and induce ACE expression in RACE+ cells.

#### Flow Cytometry

hACE transgene expression: R- and R+ mice were analyzed 7, 21, and 180 days after TMX treatment. Blood samples were collected by cardiac puncture into anticoagulant (EDTA) blood collection tubes and the spleen and one brain hemisphere were dissected after transcardiac perfusion with PBS. The spleen was homogenized with a pestle in PBS and filtered through a 70 µm strainer, and both spleen and blood samples were washed in red blood cell (RBC) lysis buffer (15 mM $NH_4Cl$, 1 mM $NaHCO_3$, and 0.11 mM EDTA) for 5 min. at RT, centrifuged to collect cell pellets and resuspended in 50 µL PBS for antibody staining. The brain tissue was chopped with scissors into fine pieces and single cells were isolated in HBSS containing 0.05% Collagenase IV, 0.1 µg/mL TLCK trypsin inhibitor, 10 µg/mL Dnase I, and 10 mM Hepes, pH 7.4 for 30 min. at 37°C. After repeated trituration, the cells were centrifuged at 500 x *g* for 10 min. at 4°C and cell pellets were mixed with 25% Percoll and centrifuged at 950 x *g* for 20 min. at 4°C. The supernatant containing cellular debris was removed and the cell pellet was resuspended in 50 µL PBS for antibody staining. Immunofluorescence labeling was performed using Fc Receptor Blocker followed by a mixture of primary antibodies: anti-CX3CR1-PE, anti-CD11b-BV605, and nuclear DNA stain, DAPI (1 mg/mL). Cells were resuspended in 500 µL MACS buffer and subjected to spectral flow cytometry analysis (Cytek, Northern Lights). Unstained and single antibody-labeled cells were used for background elimination and spectral compensation. Tissues and blood from R- mice were used as control to confirm transgene-OFF state in these animals. Data analysis was performed using FlowJo Software (BD Bioscience).

Microglia nuclei enrichment for snRNAseq: Frozen brains were quickly thawed and chopped into small pieces that were treated with Nuclei EZ lysis buffer supplemented with 0.5 U/µl RNAse inhibitor and according to manufacturer's protocol. Single nuclei were labeled with DAPI and fluorophore labeled antibodies to detect neurons (NeuN), oligodendroglia (Sox-10) and astrocytes (Sox9). To enrich for microglia, nuclei were sorted and captured on an Influx (BD Bioscience) cell sorter with capture gating set to partially exclude neurons, oligodendroglia and astrocytes (Fig. S8).

#### Microscopy:

*Wide-field and structured illumination microscopy:* Images were captured using a Zeiss AxioImager Z2 microscope equipped with Apotome 2 structured illumination optics. Monochrome fluorescent images were captured with the Zeiss Axiocam HRm camera and pseudo-colored or bright field images were captured with the Zeiss Axiocam HRc camera using Zeiss Zen software. Images were acquired using either 10x or 20x objectives.

*Laser scanning confocal microscopy:* Images were acquired on a Zeiss LSM 780 with Airyscan super-resolution optics, using a 20x or 63x oil immersion objective. Z-stack images were captured using 1 µm optical sectioning.

#### iMaris Image Analysis:

Z-stack images were reconstructed in 3-dimensions using Imaris 10.1 software. The volume of labeled structures was quantified using iMaris software to threshold mask labeled structures and reconstruct their volumes in 3-dimensions on spatially calibrated images. For all comparisons between genotypes, laser intensity and camera gain settings were identical.

Microglia Occupancy. Occupancy of relevant targets within microglia, such as $A\beta_{1-42}$, PSD-95, CD68, C1q and pSYK, were analyzed using iMaris software. Microglia were identified by Iba1 immunofluorescence signals and specific targets were identified by immunofluorescence with the appropriate primary antibody. The fluorescent signals were masked using the *Surfaces* function and thresholds were manually adjusted to accurately discriminate the overlapping volumes between the fluorescent signals. The volumes of the intersection of the labeled structure within the microglia masks were recorded using the *Overlapping Volume* or *Overlapping Volume Ratio* functions within the iMaris software.

Synapse Density. Immunofluorescence images for PSD95 and vGLUT1 were captured in the hippocampus using a 40x objective. Synapses were masked using the *Surfaces* function in iMaris software with batch surface parameters applied to entire image datasets. Thresholds were manually adjusted to accurately capture the fluorescent signals. Total puncta were measured using the *Total Count Per Image* function and PSD95-vGLUT1 colocalized pairs were quantified using the *Shortest Distance* function. Puncta pairs were classified as "colocalized" to identify a single excitatory synapse, with pre-synaptic and post-synaptic specializations, if they were within 100 nm of one another. 3-5 images were acquired per section across 3-5 mice of each genotype and sex.

Aβ Phagocytosis in Primary Microglia. Primary microglia from RACE- and RACE+ mice were isolated as described, seeded onto 8-well chamber slides and treated with HiLyte-647 labeled $A\beta_{1-42}$ that was aggregated into oligomeric form ($oA\beta_{42}$) as previously described (1). Microglia were imaged using confocal microscopy and 1 μm optical z-stack images were obtained through the entire z-plane of the cells. Immunofluorescence for CD68 and $oA\beta_{42}$ was performed and fluorescence was captured using the Imaris *Surfaces* function and the *Overlapping Volume* or *Overlapping Volume Ratio* functions to render the volume of $oA\beta_{1-42}$ that was within the CD68+ endolysosomal compartment.

Microglia Morphology. Quantification was performed by an investigator using iMaris software and blinded to the genotypes. Hippocampal Iba1-positive microglia were masked using the *Filament Tracer* algorithm and common batch parameters were applied to all groups. z-stack images with 1 μm intervals totaling 60um in depth were acquired on a confocal microscope using a 63x oil immersion objective. Individual microglia were reconstructed to quantify branch points and total process length per microglia.

Plaque Associated Microglia (PAM). Quantification was performed using Image J image analysis software with the investigator blinded to genotype. Hippocampal Iba1-positive soma within a 10 μm radius of a plaque edge were counted as plaque-associated microglia (PAM). Z-stacks with 1 μM optical section intervals were acquired using a 10x objective and projected as a single optical image to capture cells within a 10 μm radius around plaques.

#### Plaque Density and Neuron Density

Plaque density quantification was performed using ImageJ image analysis software with the investigator blinded to genotype. Images were acquired with a 10x objective to detect $A\beta_{1-42}$ immunofluorescence and for neuron density, images were acquired using a 20x objective to detect NeuN immunofluorescence. Images were converted to grayscale, subjected to auto-threshold (https://www.msn.com/en-us/feed'; *Phansalkar* method, r=10), and converted to binary. Watershed and size threshold for plaque detection was set to the value of 20 to infinity pixels for plaques and 100 to infinity pixels for neurons using the *Analyze Particle* function in ImageJ.

#### $A\beta_{1-42}$, $A\beta_{1-40}$ and Cytokine measurement

A 19-plex quantitative cytokine assay was performed as previously described (2). Briefly, micro-dissected fresh hippocampus was homogenized in RIPA buffer containing 1x protease and phosphatase inhibitor. The supernatant was utilized for cytokine and Aβ protein analyses and immunoblotting. All samples were assayed in duplicate using the MSD V-PLEX Mouse Cytokine Assay kit and the MSD V-PLEX Aβ Peptide Panel (6E10) kit to quantify levels of interferon gamma (IFN-γ), interleukin (IL)-1β, IL-2, IL-4, IL-6, IL-10, IL-12p70, C-X-C motif chemokine ligand 1 (CXCL1), tumor necrosis factor alpha (TNF-α), $A\beta_{38}$, $A\beta_{40}$ and $A\beta_{42}$.

#### Tissue Microarray

Brains from 3 male and 3 female 6 month-old mice with genotypes R-;AD-, R+;AD-, R-;AD+ and R+;AD+ were formalin fixed and paraffin embedded. Paraffin tissue blocks were faced until matching anatomical regions were visible across all blocks and 3 hippocampal donor cores (1mm diameter) were extracted from each block and placed into a recipient tissue microarray block using a GrandMaster TMA (3D Histotechnology, Hungary) instrument.

#### GeoMx Digital Spatial Profiling (Nanostring)

The TMA block was faced until all tissue cores were represented in full circumference and sections of 4 µm thickness were used for analysis. Sections were labeled by immunofluorescence with direct fluorophore conjugated morphology markers to detect microglia (Iba1-Alexa6470), Aβ plaques (6E10-Alexa488) and nuclei (SYTO Green). The slides were sequentially hybridized using RNA probes from the Mouse Whole Transcriptome Atlas library kit according to the manufacturer's protocol. The morphology markers were used to localize Aβ plaques and microglia to draw regions of interest (ROIs) for gene expression sampling using the GeoMx Digital Spatial Profiler Instrument software. ROIs that contained many plaque-associated microglia (PAM) and ROIs that contained many non-plaque-associated microglia (nPAM) were selected and the microglia fluorescence channel (Iba1) was used as the ultraviolet (UV) light mask to photocleave the hybridized nucleic acid probes that were preferentially bound to microglia in the ROIs. Photocleaved probe libraries were generated and Illumina NextSeq500 sequencing was performed to interrogate gene expression in the microglia within the ROIs. Differential gene expression analysis was performed using the Nanostring GeoMx DSP platform control and analysis software and integrated Reactome Pathway Analysis.

#### snRNAseq (10x Genomics)

Sorted nuclei enriched for microglia nuclei from 6-month-old male and female R-;AD-, R+;AD-, R-;AD+ and R+;AD+ mice were divided into two groups so that nuclei from all genotypes and sexes could be processed efficiently using hashing antibodies to multiplex the samples by genotype and sex. Six hash tagging antibodies were used to label groups of nuclei as described in manufacturer's protocols (Biolegend). Hash tag labeled nuclei were captured using a Chromium Chip G microfluidics chamber on a Chromium X series processor using the Chromium Next GEM 3' kit as specified by the manufacturer (10X Genomics). cDNA libraries were processed for compatibility with Illumina sequencing using the 3' Feature Barcode Kit, the Dual Index Kit NT Set A, and the Dual Index Kit TT Set A according to the manufacturer's specifications (10X Genomics).

Sequenced samples were demultiplexed and aligned to the GRCm38 (mm10) reference sequence dataset using the Cellranger *Count* pipeline with the *include-introns* flag set. Notably, the microglia expressed DAM gene *Clec7a* was not represented in the GRCm38 reference dataset. The Seurat toolkit (version 4.3.0), implemented in the R language, was used for downstream analysis (3). The UMI count matrices were filtered to exclude nuclei with less than 200 genes and more than 5000 genes. Counts were normalized using the CLR method prior to running *DoubletFinder* (version 2.0.3) assuming an 8% doublet rate (4). Subsequently, the *FindVariableFeatures()* function was implemented using a variance stabilizing transformation to identify the 2,000 most variable genes. Data from two batches of samples were subsequently integrated with the *FindIntegrationAnchors()* function, using default parameters, and the *IntegrateData()* function with 16 principal components (PC). The final dataset consisted of 16,466 nuclei, with a mean UMI count of 4,664 counts per nucleus and 2,062 genes per nucleus. Integrated data were scaled using the *ScaleData()* function with the linear model, prior to dimension reduction with *RunPCA()*. Clusters were identified with the *FindNeighbors()* function constructing a KNN graph based on the Euclidean distance in PCA space (based on the first 16 PC) with edge weight between cells based on their Jaccard similarity. Following this, modularity optimization with the Louvain algorithm, to iteratively group cells together, was performed using the *FindClusters()* function with a resolution of 0.4. The *Uniform Manifold Approximation* and *Projection* embeddings were then calculated and plotted. The Microglia clusters, constituting 1,588 nuclei, were subsequently subset out and re-clustered according to the outline above.

Cluster analysis. Markers for all clusters were obtained using the *Findmarkers()* function with default settings, selecting only genes with positive fold change. Differential expression analysis to compare between genotypes and between microglial subsets was also conducted using the *Findmarkers()* function. Pseudobulking gene expression by genotype and subset was conducted with the *AverageExpression()* function.

#### Behavioral Tests

Locomotor activity and open field testing. Exploratory activity and anxiety-like behavior were measured using an open-field apparatus (30 x 30 x 30 cm). The center zone was defined as a square, 10 cm away from the wall. Each mouse was placed in the center of the apparatus and allowed to freely explore the area for 10 min. with video camera tracking. Anxiety-like behavior was assessed by calculating time spent in the center relative to the perimeter. The behaviors were analyzed by ANY-maze software.

Barnes maze (5). Barnes Maze testing was performed to assess learning and memory using a round table with 20 holes equidistant from the center. During testing, an escape box was placed under one of these holes, while false boxes too small to be entered, were placed beneath the other 19 holes. The apparatus was brightly illuminated in a dark room with three overhead LED lamps (~800 lux). Mice were trained to locate a dark escape box hidden underneath a hole positioned around the perimeter of the apparatus with two trials per day for five consecutive days. The trial was initiated by placing the animal in the center of the maze covered under a cylindrical starting chamber. After 10 sec. delay, the starting chamber was lifted to allow the animal to freely escape. The training session ended after the animal entered the escape box or when 5 min. had elapsed. The animal remained in the escape box for an additional 30 sec. before being removed and transferred to the home cage. All boxes and the maze surface were sprayed with 70% ethyl alcohol and then wiped to remove odor cues for each subsequent trial. The location of the escape box remained the same during every trial of the training/acquisition phase. A trial was given two days following the last learning day to assess memory retention. Following the memory retention phase, a reversal phase of training was begun. In reversal phase, the escape box was placed in a rotated location 180° from the initial training box. Using the same procedure with training/acquisition phase, reversal trials were repeated two times per day over three consecutive days. The behaviors were analyzed using ANY-maze software (Stoelting Co. USA).

Contextual and cued fear conditioning (6). Fear conditioning was conducted in a purpose-built fear conditioning chamber (Maze Engineers). The chamber consisted of a Plexiglas box with an electrified floor grid placed inside of a sound-proof cubicle (55 cm W × 48 cm D × 551 cm H). ANY-maze software was used to control the delivery of tones and foot shocks. For three days prior to fear conditioning training, animals were habituated to handling. During animal training and testing, the technician remained blinded to genotypes. On training and testing days, animals were transferred in their home cages to a room contiguous with the training room where they were allowed to acclimate for 1 hour. For training, animals were transferred to the fear conditioning chamber and left undisturbed for 180 sec., followed by 5 tone-shock pairings separated by a 1min. interval. Each tone-shock pair consisted of a 30 sec. 2800 Hz, 85 dB tone and a 2-sec. 0.75 mA shock delivered during the last 2 sec. of the tone and animal freezing was measured for 1 min. which was detected automatically with ANY-maze software. Contextual fear memory was tested 24 hours after training and cued fear conditioning memory was tested 36 hours after training.

Contextual fear memory testing was performed by returning the animals to the same training chamber and freezing was measured for 5 minutes without applying a shock. Freezing was detected automatically with ANY-maze software using a minimum freezing duration of 2 milli-sec. Fear memory was calculated as the percent of time that animals spent freezing during the total time in chamber.

Cued fear memory testing was performed 12 hours after the contextual fear memory using a different chamber environment context. The test consisted of a pre-tone period of 180 sec., followed by the two 30-sec. 2800 Hz, 85 dB tones for 1 min. each without any foot shock. Cued fear conditioning memory was measured for 5 minutes after the tone stimuli and fear memory was calculated as the percent of time that animals spent freezing during the 5 min. measurement period.

**Uptake and endolysosomal trafficking assay using Incucyte:**

oA$\beta_{1-42}$-pHrodo conjugation: A$\beta_{1-42}$ peptide was aggregated into oligomeric (oA$\beta_{1-42}$) form and then conjugated to succinimidyl pHrodo-Red using manufacturer's protocols (Invitrogen).

Primary microglia were isolated from R-;AD- (R-) and R+;AD- (R+) mice, seeded in 96 well plates (3.5x$10^4$ cells per well) and incubated at 37 °C under 5% $CO_2$ for 24hrs. Two days prior to experiments the microglia were treated with 4-OHT and then incubated on ice for 10 min. while Zymosan-pHrodo bioparticles or oA$\beta_{1-42}$-pHrodo was added to the medium. Cells were then imaged using the phase contrast channel and the red channel on the IncuCyte SX5 platform (Sartorius, Germany). Nine images from distinct regions within each well were taken at intervals of 30 min. over 5 days using a 20x objective. The images were analyzed using the IncuCyte SX5 Basic Software. The red channel acquisition time was 400ms and cell segmentation was performed using the phase contrast channel to match to the fluorescent channel signals. For uptake and trafficking measurements, curve integration was performed and measurements were normalized to signals obtained from wild type cells.

**Ratiometric analysis:**

Genes identified as differentially expressed in plaque-associated microglia (PAM) by Nanostring GeoMx spatial transcriptomics were analyzed in the dataset generated by snRNA-seq. To assess whether the DAM-A snRNA-seq microglial cluster had increased expression of genes identified by spatial transcriptomics as expressed in PAM, average UMI counts were compared between DAM-A and DAM-B, and displayed as a ratio between DAM-A and DAM-B counts.

**Statistics overview:**

Statistical analyses were performed in GraphPad Prism using the tests shown for each figure in the table below. $p < 0.05$ was considered statistically significant and p values were abbreviated in all figures as: * = $p < 0.05$, ** = $p < 0.01$, *** = $p < 0.001$ and **** = $p < 0.0001$.

Results for males and females were pooled together since no individual sex differences were identified.

| Figure | Test Type | Post Hoc. | M/F Differences | Notes |
|---|---|---|---|---|
| 2C | Two Way ANOVA | Tukey | No | |
| 2D | Mann Whitney | | No | |
| 3A | Mann Whitney | | No | |
| 3C | Unpaired T test | | No | |
| 3D | Welsh's T test | | No | |

| Figure | Test Type | Post Hoc. | M/F Differences | Notes |
|---|---|---|---|---|
| 3F | Unpaired T test | | No | |
| 3G | Unpaired T test | | No | |
| 3H | Kruskall-Wallis | Dunn | No | |
| 4A | One Way ANOVA | Tukey | No | |
| 4B | Two Way ANOVA | Tukey | No | |
| 4C | Kruskall-Wallis | Dunn | No | |
| 4D | Kruskall-Wallis | Dunn | No | |
| 5A | Two Way ANOVA | Tukey | No | Outlier Detection, Q=0.1% |
| 5B | One Way ANOVA | Tukey | No | Outlier Detection, Q=0.1% |
| 5C | Brown-Forsythe and Welsh ANOVA | Tamhane T2 | No | Outlier Detection, Q=0.1% |
| 5D | Two Way ANOVA | Tukey | No | Outlier Detection, Q=0.1% |
| 5E | One Way ANOVA | Tukey | No | Outlier Detection, Q=0.1% |
| 5F | One Way ANOVA | Tukey | No | Outlier Detection, Q=0.1% |
| 8A | Kruskal-Wallis | Dunn | No | |
| 8B | Brown-Forsythe and Welsh ANOVA | Tamhane T2 | No | |
| 8C | Unpaired T-test (RT-PCR results)<br><br>One-way ANOVA | <br><br><br>Tukey | No | |
| S3 | Brown-Forsythe and Welsh ANOVA | Tamhane T2 | No | |
| S4 | Two Way ANOVA | Tukey | No | |
| S5 | Unpaired T-test | | No | |
| S6A | Kruskall-Wallis | Dunn | No | |
| S6B | Kruskall-Wallis | Dunn | No | |
| S6C | Kruskall-Wallis | Dunn | No | |
| S6D | Kruskall-Wallis | Dunn | No | |
| S6E | Kruskall-Wallis | Dunn | No | |
| S10B | Unpaired T test | | N | |

**References.**

1. He Y, Wei M, Wu Y, Qin H, Li W, Ma X, et al. Amyloid β oligomers suppress excitatory transmitter release via presynaptic depletion of phosphatidylinositol-4,5-bisphosphate. *Nature communications.* 2019;10(1):1193.

2. Oblak AL, Lin PB, Kotredes KP, Pandey RS, Garceau D, Williams HM, et al. Comprehensive Evaluation of the 5XFAD Mouse Model for Preclinical Testing Applications: A MODEL-AD Study. *Front Aging Neurosci.* 2021;13:713726.

3. Butler A, Hoffman P, Smibert P, Papalexi E, and Satija R. Integrating single-cell transcriptomic data across different conditions, technologies, and species. *Nature biotechnology.* 2018;36(5):411-20.

4. McGinnis CS, Murrow LM, and Gartner ZJ. DoubletFinder: Doublet Detection in Single-Cell RNA Sequencing Data Using Artificial Nearest Neighbors. *Cell Syst.* 2019;8(4):329-37.e4.

5. Tajti BT, Yoon O, Ernyey AJ, Gáspár A, Varga BT, and Gyertyán I. Using Appetitive Motivation to Train Mice for Spatial Learning in the Barnes Maze. *Biomed Res Int.* 2023;2023:6625491.

6. Wehner JM, and Radcliffe RA. Cued and Contextual Fear Conditioning in Mice. *Current Protocols in Neuroscience.* 2004;27(1):8.5C.1-8.5C.14.

7. Keren-Shaul H, Spinrad A, Weiner A, Matcovitch-Natan O, Dvir-Szternfeld R, Ulland TK, et al. A Unique Microglia Type Associated with Restricting Development of Alzheimer's Disease. *Cell.* 2017;169(7):1276-90 e17.

#### Extended Results:

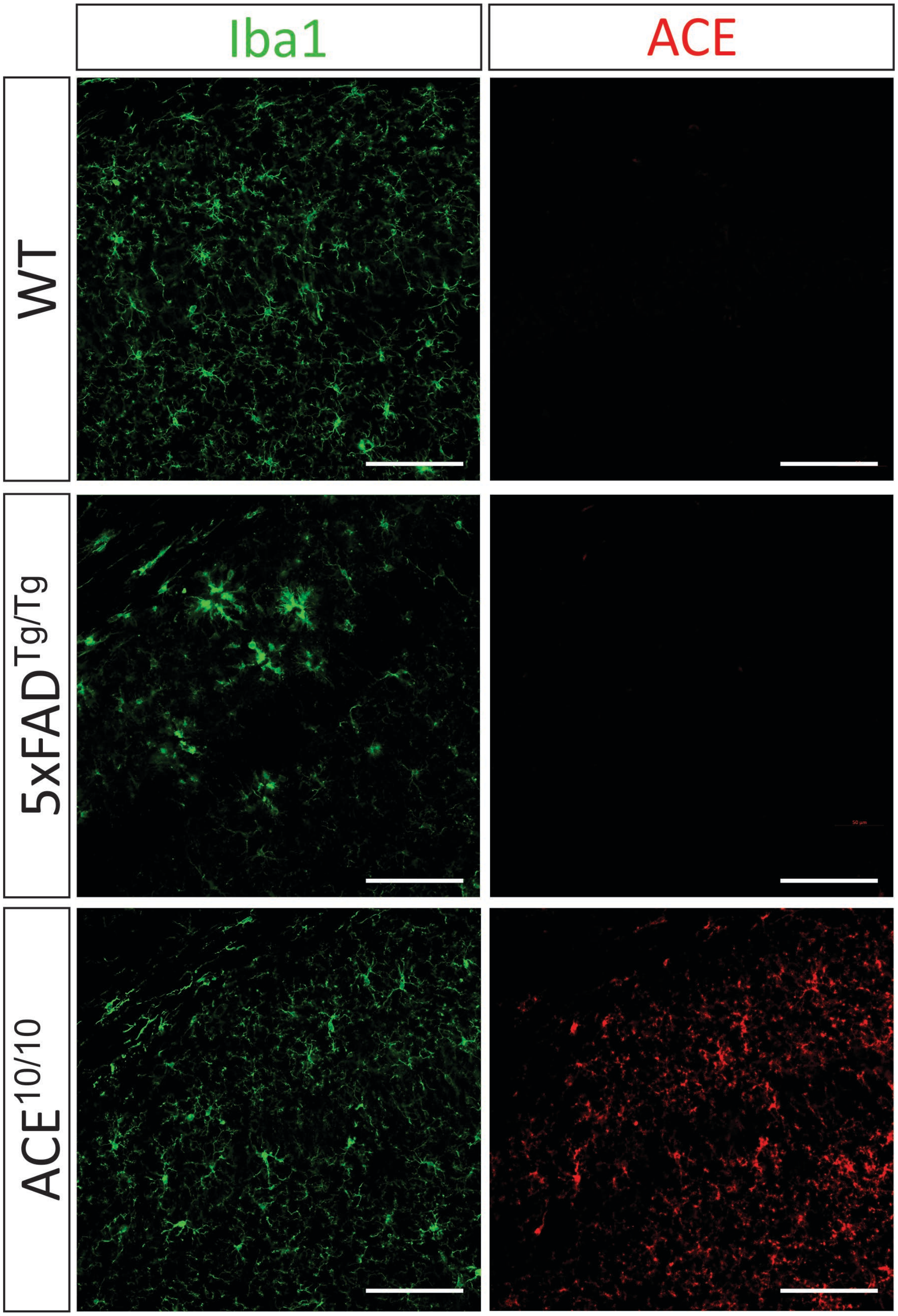

**Fig. S1: Microglia express ACE in $ACE^{10/10}$ mice**. ACE expression is detected in Iba1 positive microglia by immunofluorescence in $ACE^{10/10}$, but not wild type (WT) or 5xFAD mice. (scale = 100 µm)

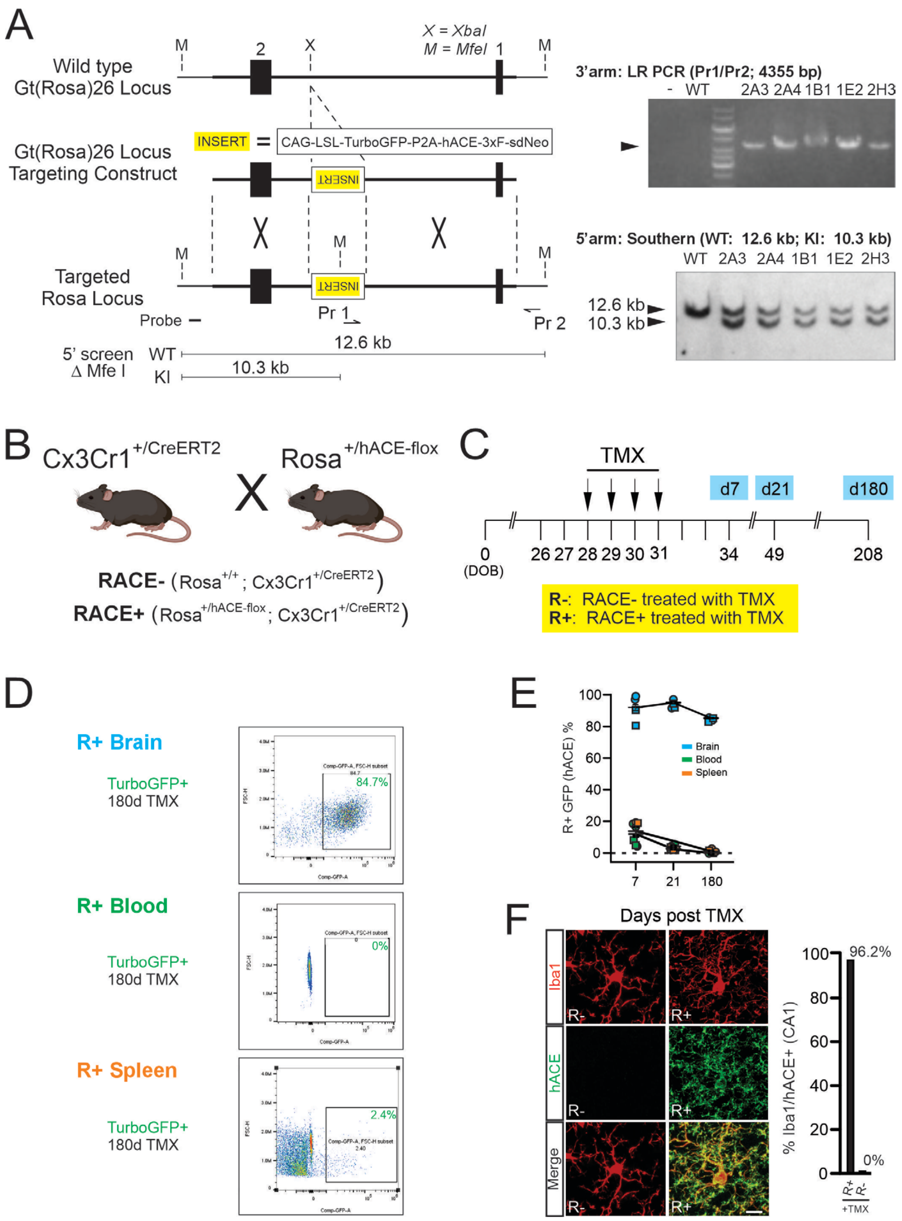

**Fig. S2: ACE expression in microglia in vivo.** **(A)** Targeting construct to conditionally express TurboGFP and flag-tagged human ACE (hACE-3xF). Clones were screened by PCR across the 3' homology domain and by Southern blotting across the 5' homology domain. Targeted clone 2A3 was used to generate 3 male and 2 female germline heterozygous ACE-flox (Rosa$^{+/hACE\text{-}flox}$) mice. **(B)** ACE-flox mice were mated to Cx3Cr1$^{+/CreERT2}$ mice to generate RACE- (Cx3Cr1$^{+/CreERT2}$; Rosa$^{+/+}$) and RACE+ (Cx3Cr1$^{+/CreERT2}$; Rosa$^{+/hACE\text{-}flox}$) mice. **(C)** RACE- and RACE+ mice were injected with Tamoxifen (TMX; 125 mg/kg, IP) to activate Cre-recombinase in R- and R+ myeloid-derived cells. **(D)** Myeloid cells isolated from brain, spleen and blood were gated on DAPI, CD11b and Cx3Cr1 and then sub-gated on GFP to detect transgene expression, 7, 21 and 180 days after TMX injection. **(E)** Transgene expression persisted in myeloid cells in the brain and became undetectable in the blood and spleen by 3 weeks after TMX injection. **(F)** >96% of microglia in the brain of R+ mice expressed high levels of ACE that was not detectable in microglia from R- mice. (Scale bar = 10 μM).

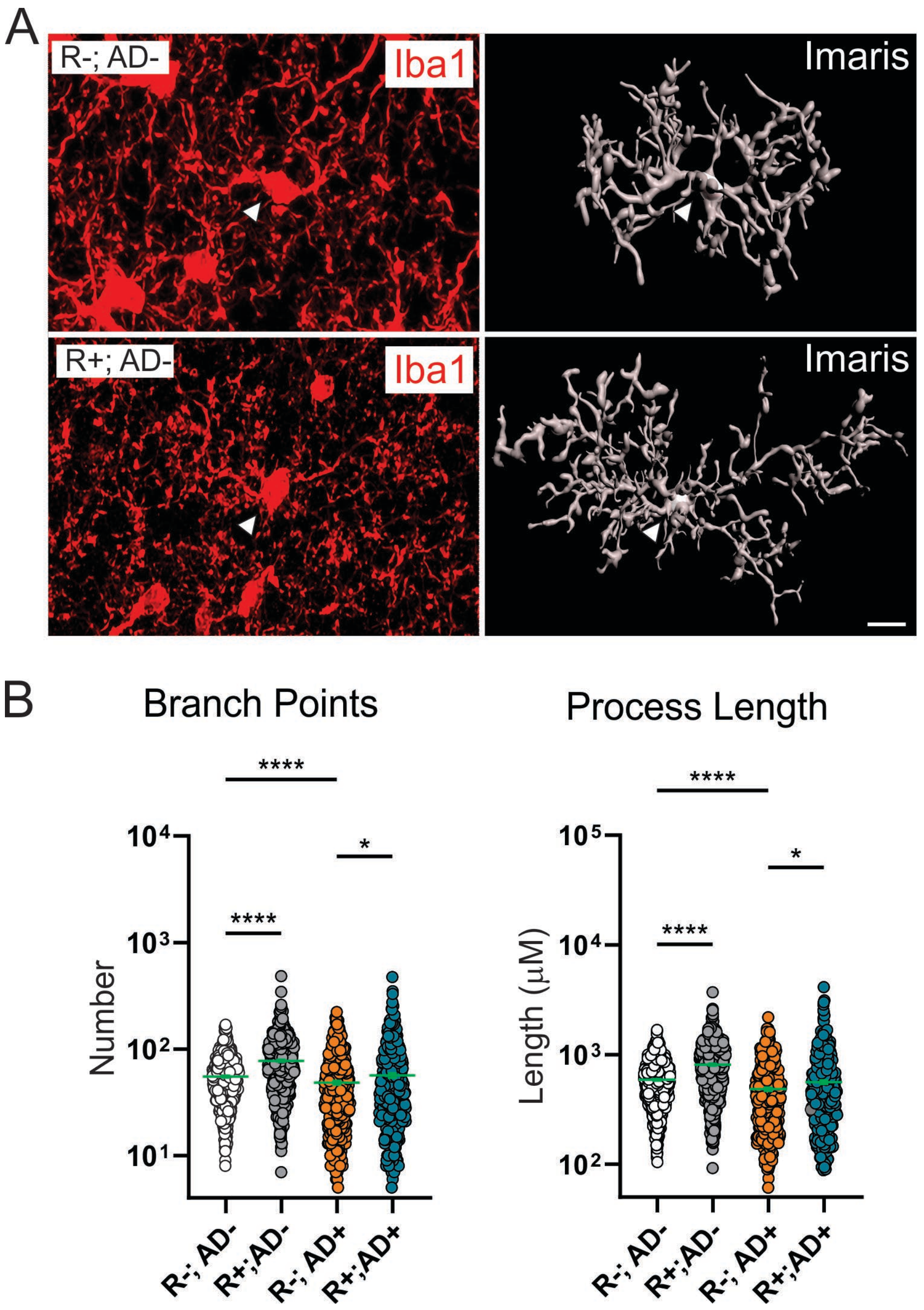

**Fig. S3: Changes in morphology mediated by Aβ deposition and ACE expression in microglia**. **(A)** Microglia cell bodies and their processes in CA1 were labeled with Iba1 and reconstructed using iMaris software filament tracing on 50 μM thick tissue sections (scale = 10 μM). **(B)** Process branching and length were increased in R+;AD- (N=738) relative to R-;AD- (N= 710) microglia ($p < 0.0001$). They were decreased in R-;AD+ microglia (N=413) compared to R-;AD- microglia ($p< 0.0001$), consistent with morphological changes associated with immune activation in response to Aβ. In ACE expressing microglia from R+;AD+ mice (N=448), process branching and length was restored to levels observed in wild type (R-;AD-) microglia.

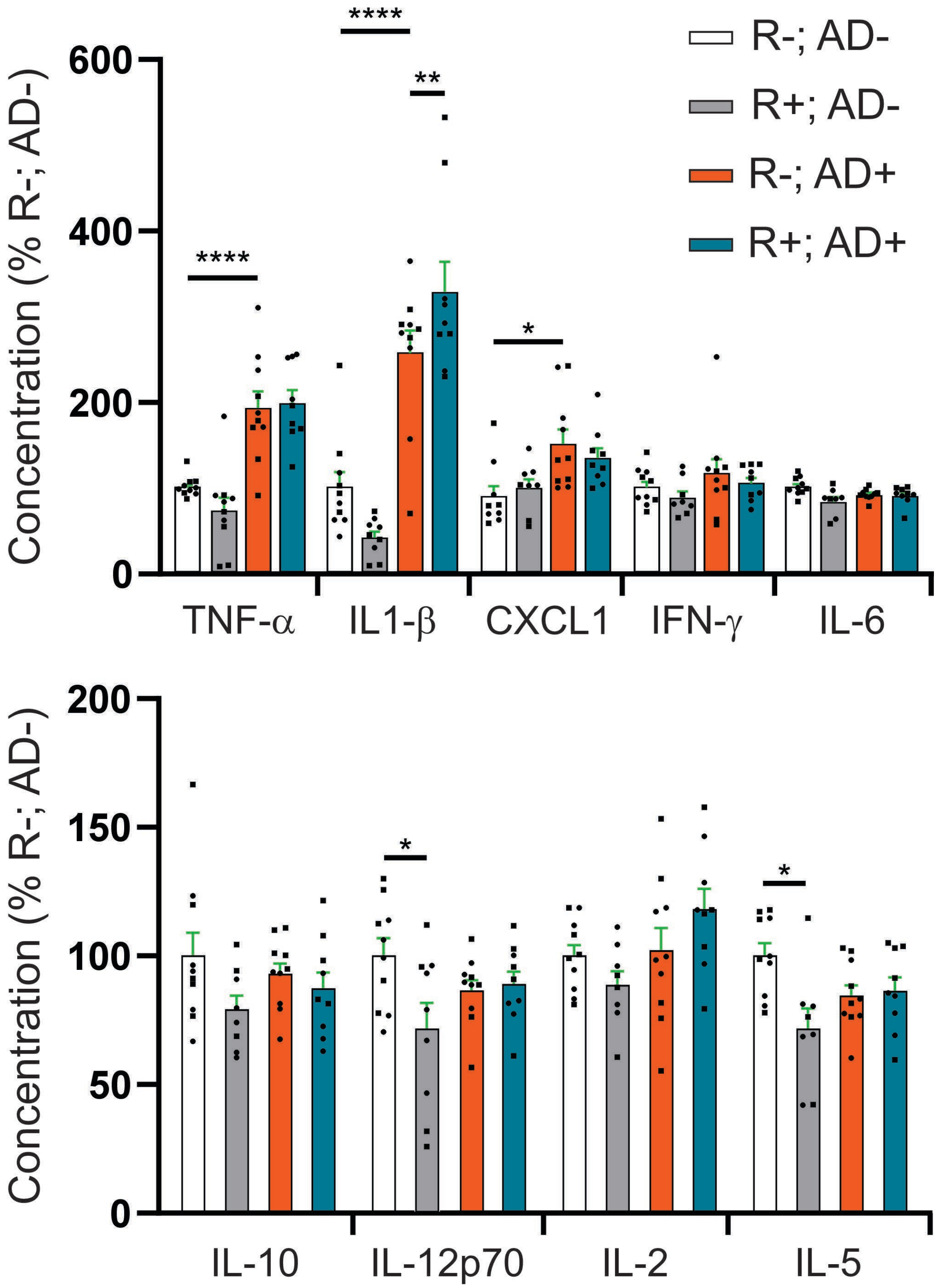

**Fig. S4: Cytokine levels in the brains of transgenic mice are minimally affected by ACE expression in microglia.** Of nine cytokines measured from forebrain protein lysates, only TNF-α and IL1-β were elevated in R-;AD+ compared to R-;AD- brains. ACE expression in microglia from R+;AD+ mice slightly increased IL1-β levels further in R+;AD+ compared to R-;AD+ brains ($p < 0.01$) and it decreased IL-12p70 and IL-5 compared to all other genotypes. (male mice = round symbol and female mice = square symbol)

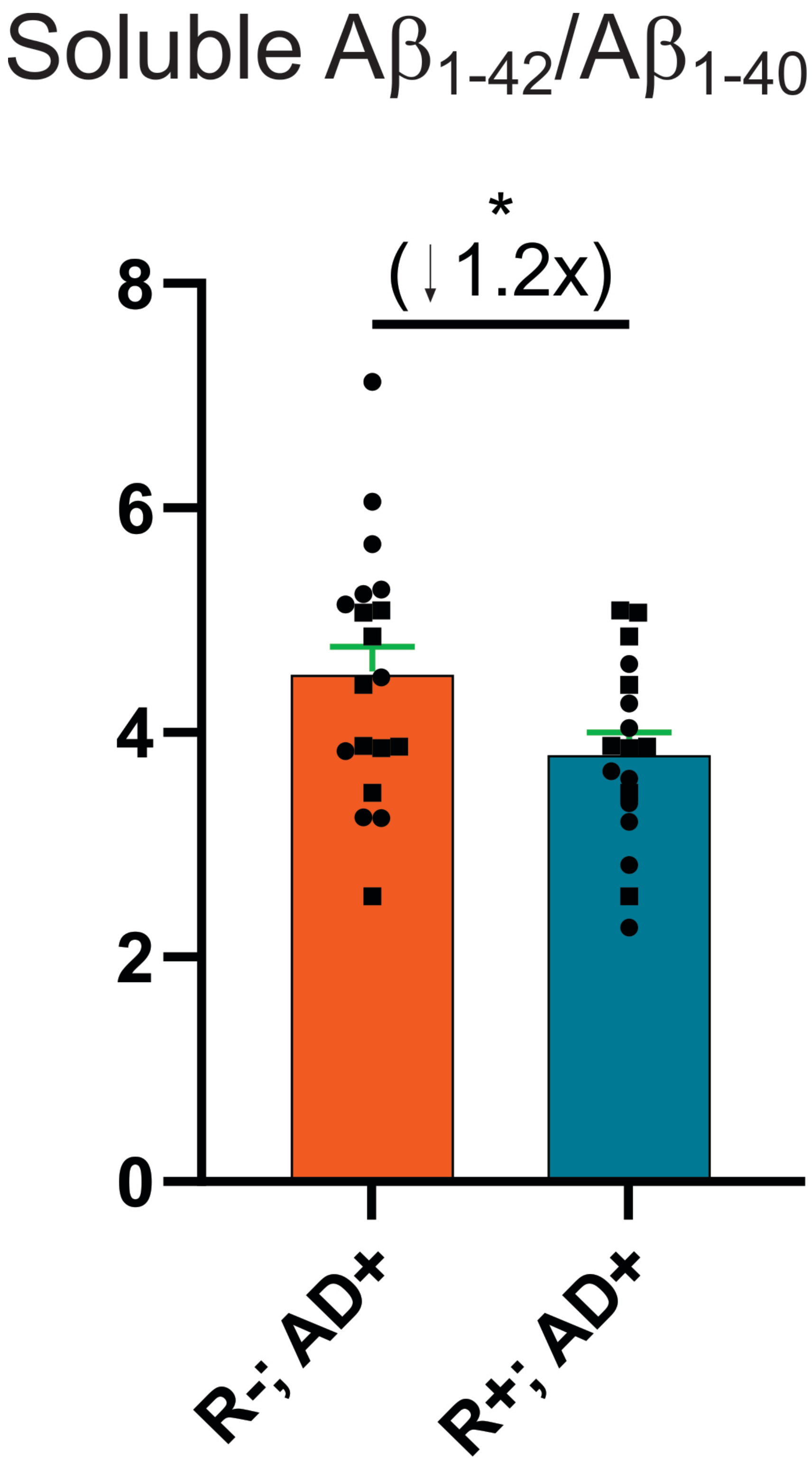

**Fig. S5: $A\beta_{1\text{-}42}$ to $A\beta_{1\text{-}40}$ ratio in brains of transgenic mice is slightly decreased in microglia expressing ACE**. ACE expression in microglia slightly reduces the ratio of $A\beta_{1\text{-}42}$ to $A\beta_{1\text{-}40}$ by 1.2-fold in R+;AD+ compared to R-; AD+ mice ($p < 0.05$). (male mice = round symbol and female mice = square symbol)

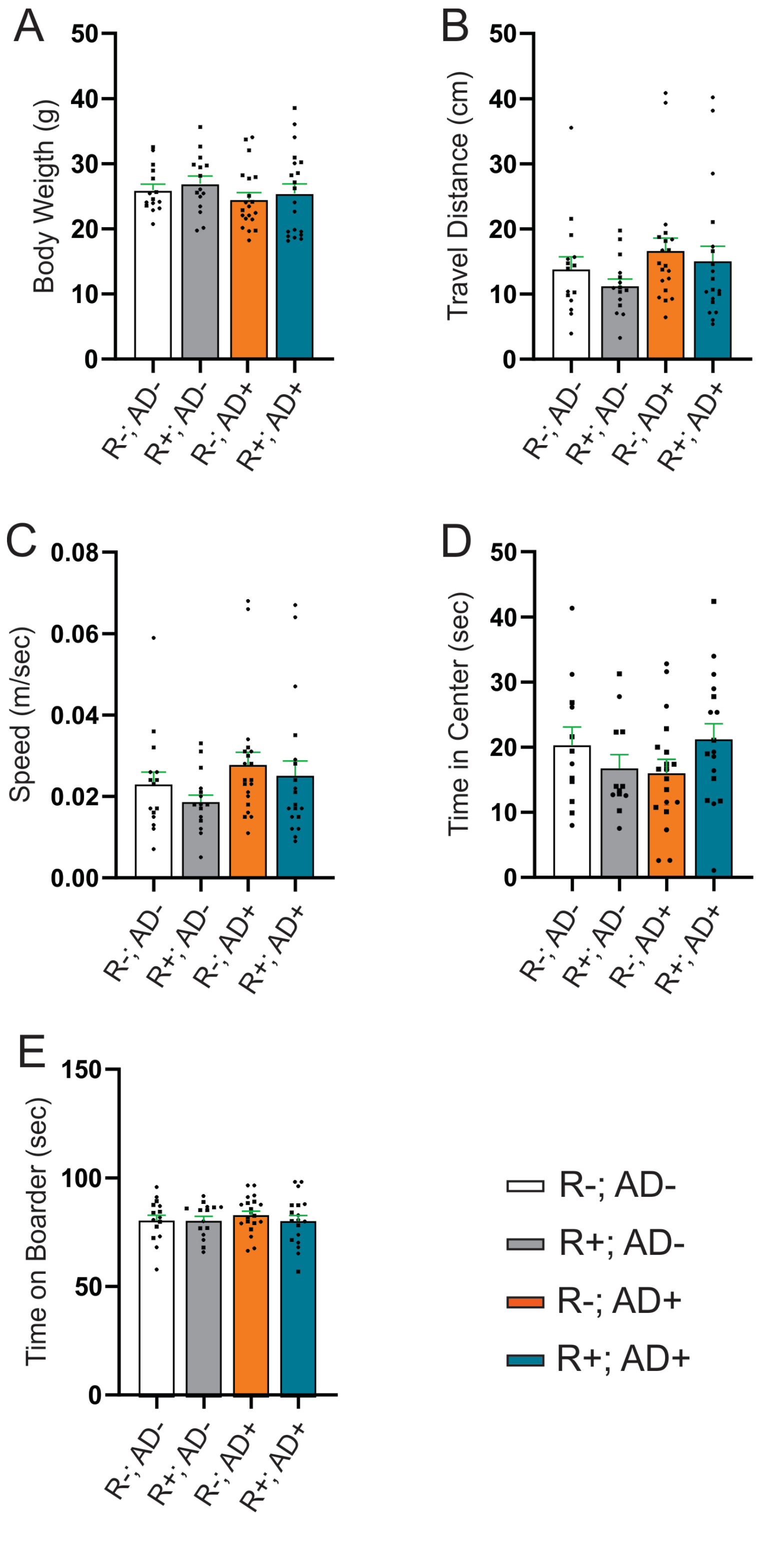

**Fig. S6: Normal body weight, locomotor activity and anxiety behavior in R+;AD-, R-;AD+ and R+;AD+ compared to R-;AD- mice.** **(A)** Body weight, **(B)** locomotor distance, **(C)** locomotor speed, **(D)** time spent in center and **(E)** time spent on boarder in open field testing were similar between all genotypes. (male mice = round symbol and female mice = square symbol)

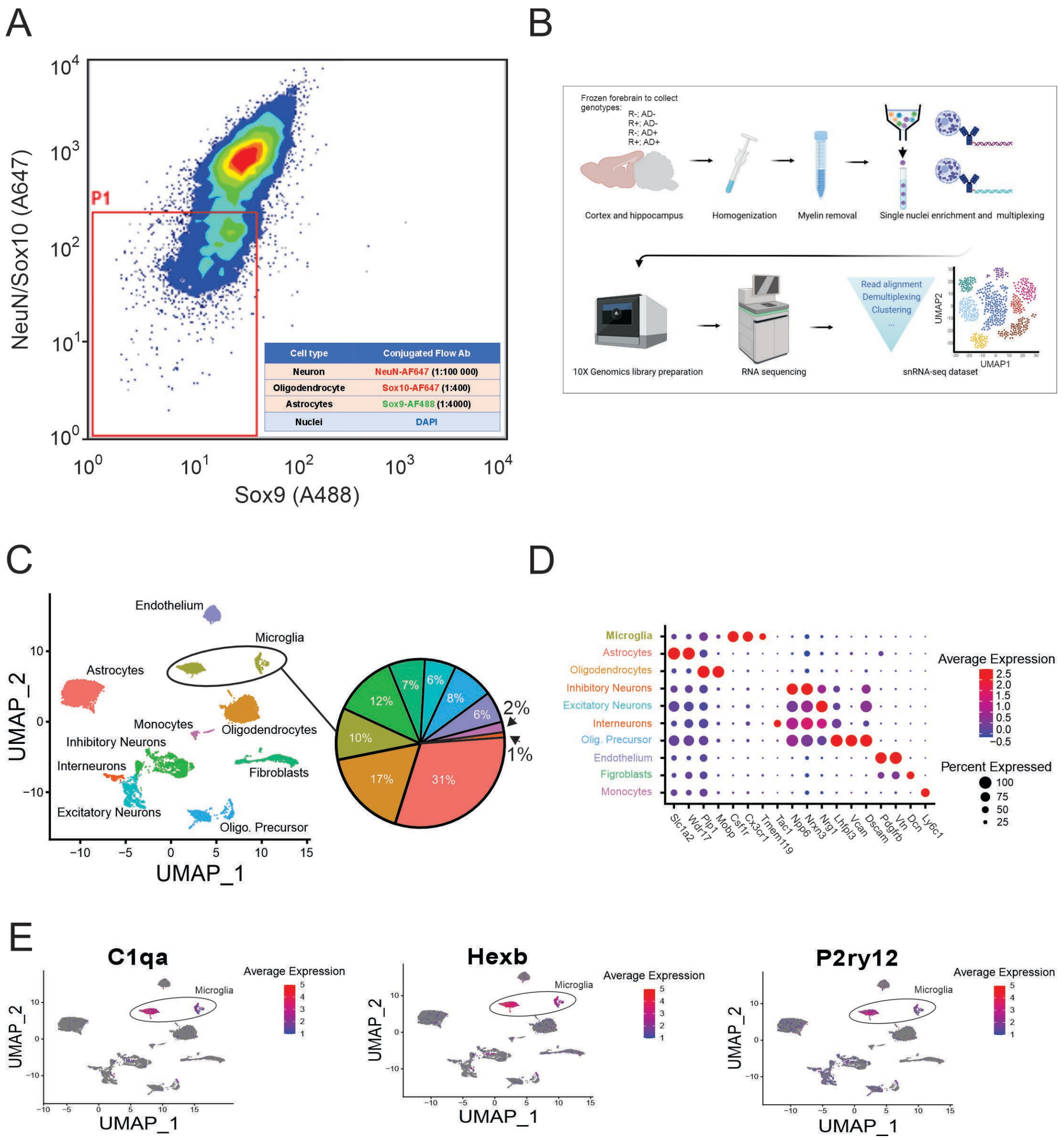

**Fig. S7: snRNAseq identifies molecular changes modulated by ACE expression in microglia. (A)** Frozen forebrains from 6-month-old R-;AD-, R+;AD-, R-;AD+ and R+;AD+ mice were dissociated into single nuclei and enriched for microglia nuclei using fluorescence activated cell sorting (FACS) to diminish neuronal nuclei that express NeuN, oligodendroglia nuclei that express Sox10 and astrocyte nuclei that express Sox9. To maximize microglia nuclear enrichment, the capture gate (P1) was set to also include some NeuN+, Sox10+ and Sox9+ nuclei. **(B)** Single nuclei were labeled with multiplexing antibodies to trace their genotypes and they were captured using the 10x Genomics barcode Gel Bead system. Transcriptomes within single nuclei were reverse transcribed and cDNA libraries were prepared for sequencing, genotype demultiplexing and cluster analysis. **(C)** Single nuclei were organized into UMAP clusters and the relative percent of nuclei within each cluster identified. Microglia were enriched from their normal 2-3% frequency to 10% frequency. **(D)** Well-established cell lineage-dependent genes were used to assign cell lineage identity to each cluster. **(E)** C1qa, Hexb and P2ry12, genes expressed relatively specifically in the two clusters identified as microglia (all cell nuclei shown in grey).

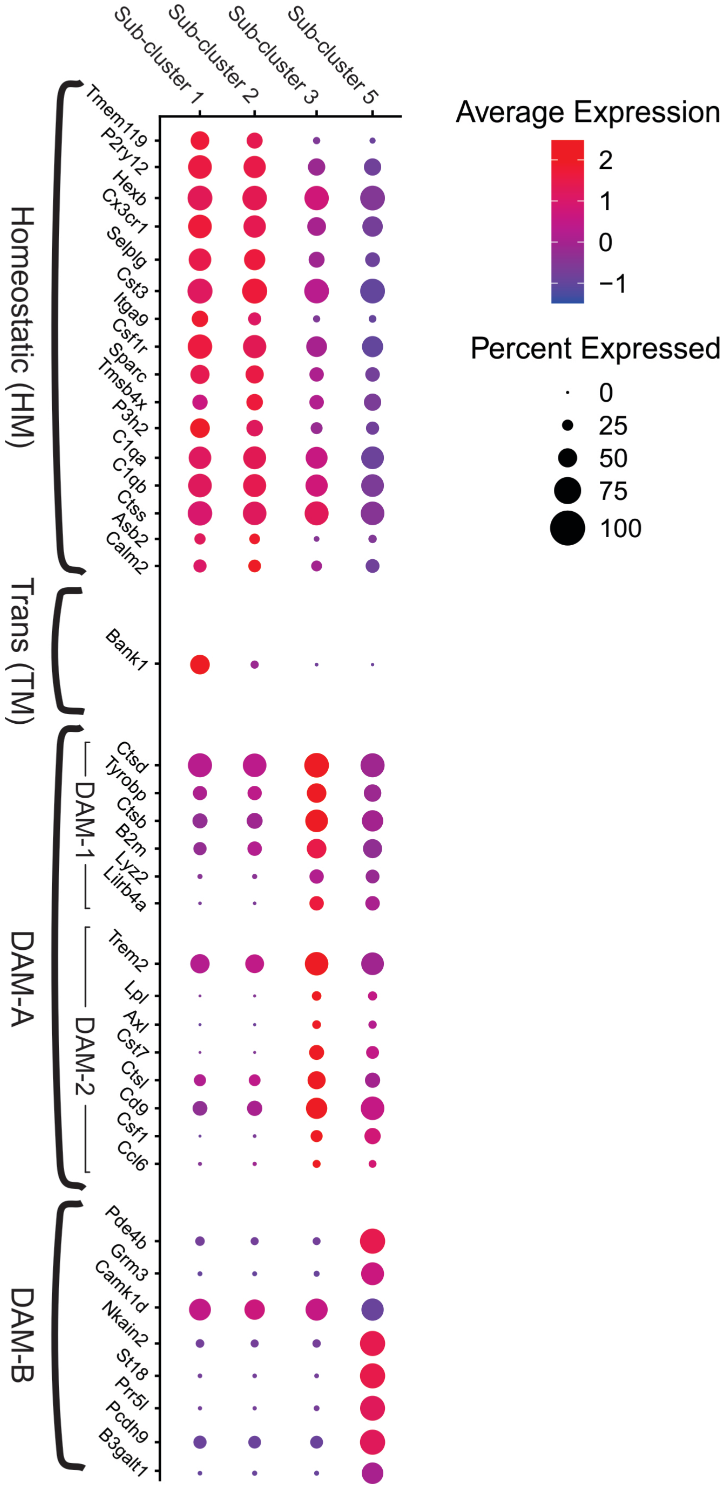

**Fig. S8: Microglial subclusters are identified by gene expression analysis**. Subclusters were identified as homeostatic microglia (HM; subcluster 1), transitional microglia (TM; subcluster 2) and disease associated microglia (DAM; subclusters 3 and 5) by gene expression analysis. The DAM-A (subcluster 3) contains the previously characterized TREM2-independent DAM stage 1 and TREM2-dependent DAM stage 2 microglia (7).

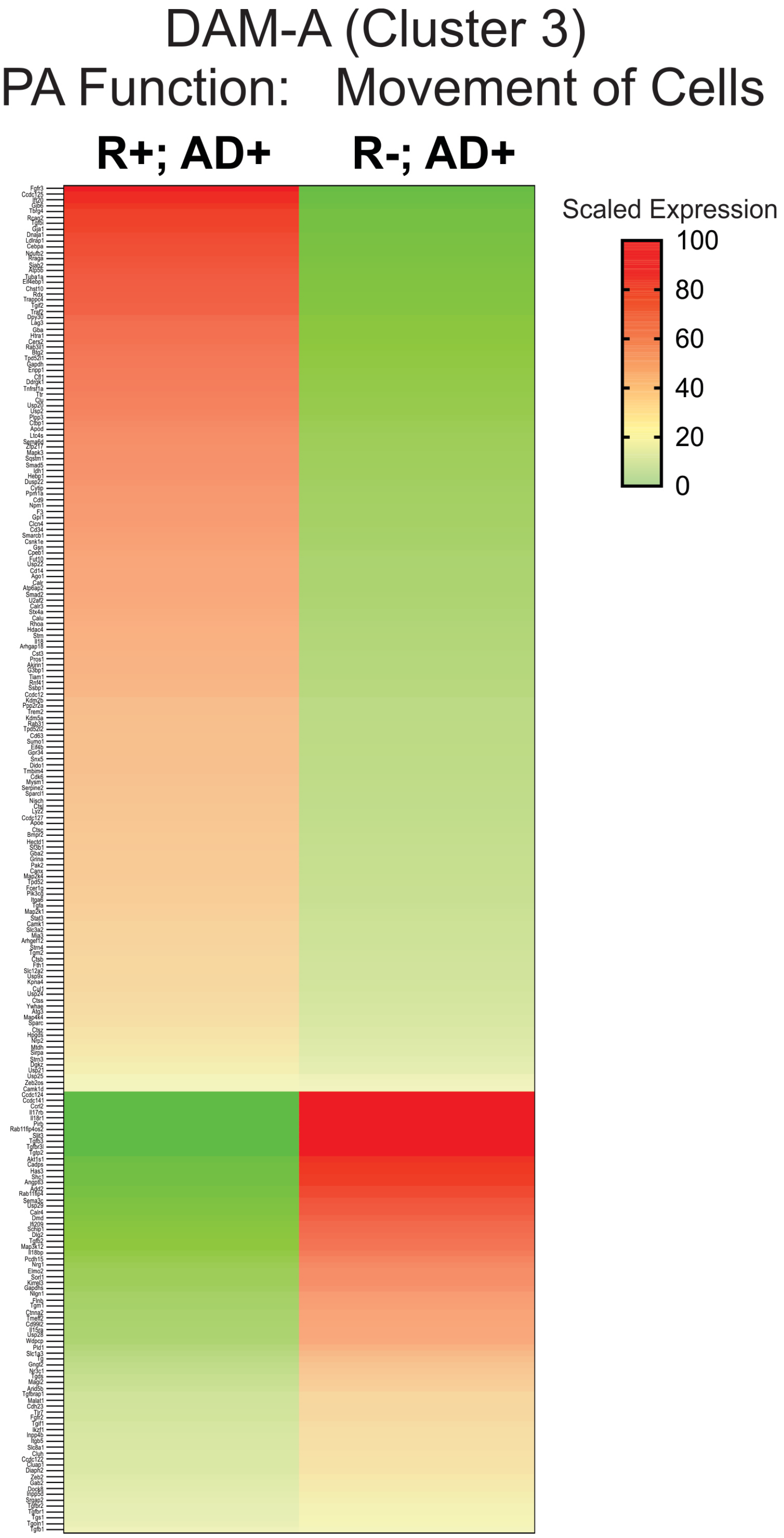

**Fig. S9: ACE expression in microglia modulates gene expression associated with cellular mobility in DAM-A microglia.** IPA function for *Movement of Cells* was significantly altered by ACE expression in DAM-A microglia. Many genes in the migration pathway were significantly modulated by ACE expression.

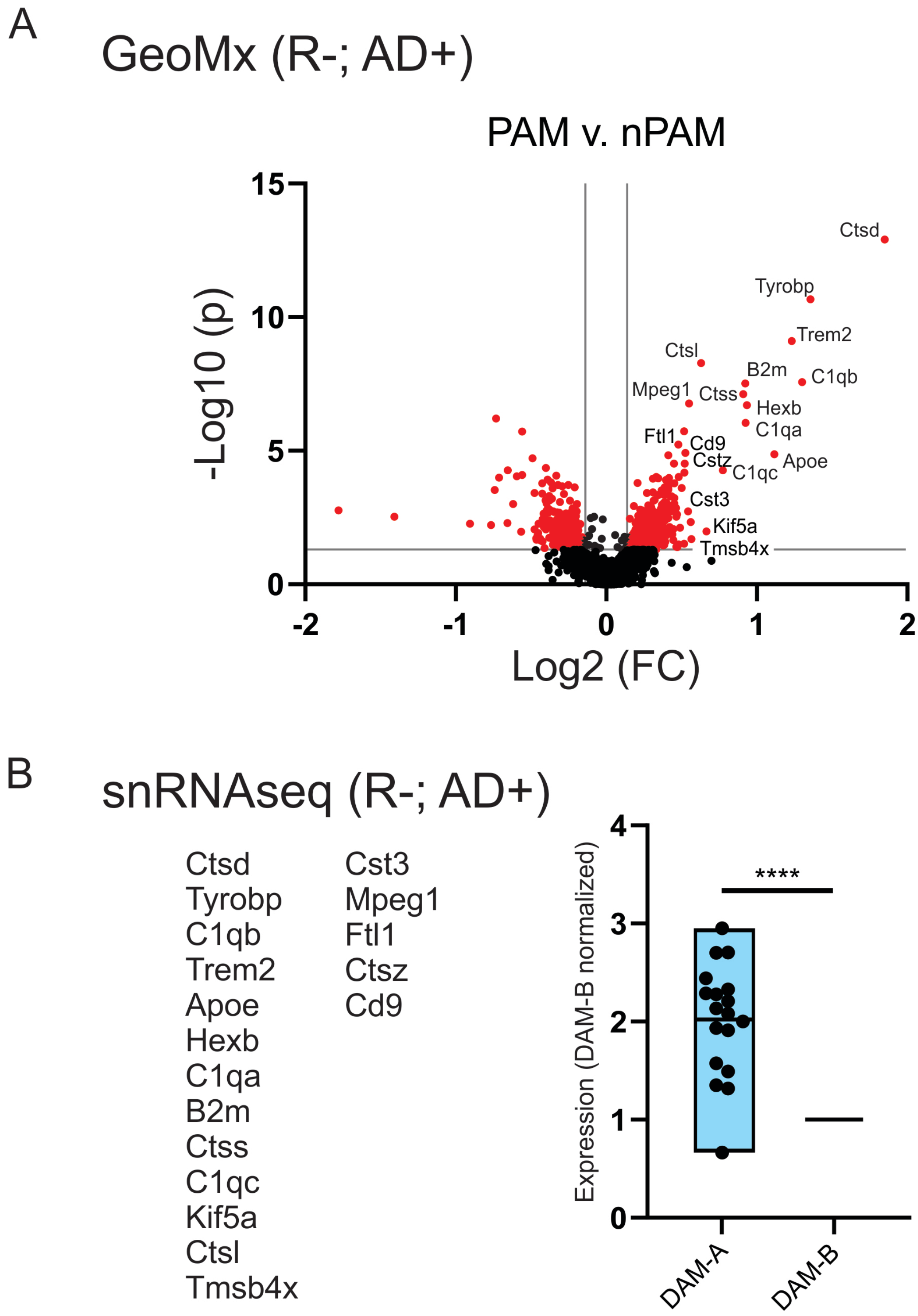

**Fig. S10: Spatial transcriptomics identifies differentially expressed genes in plaque-associated microglia (PAM) that are preferentially localized to DAM-A microglia. (A)** Volcano plot shows genes differentially regulated in PAM relative to non-plaque-associated microglia (nPAM) using Nanostring GeoMx. **(B)** The most highly differentially upregulated genes in PAM on tissue sections (Nanostring GeoMx) are associated with DAM-A microglia in snRNAseq analysis.

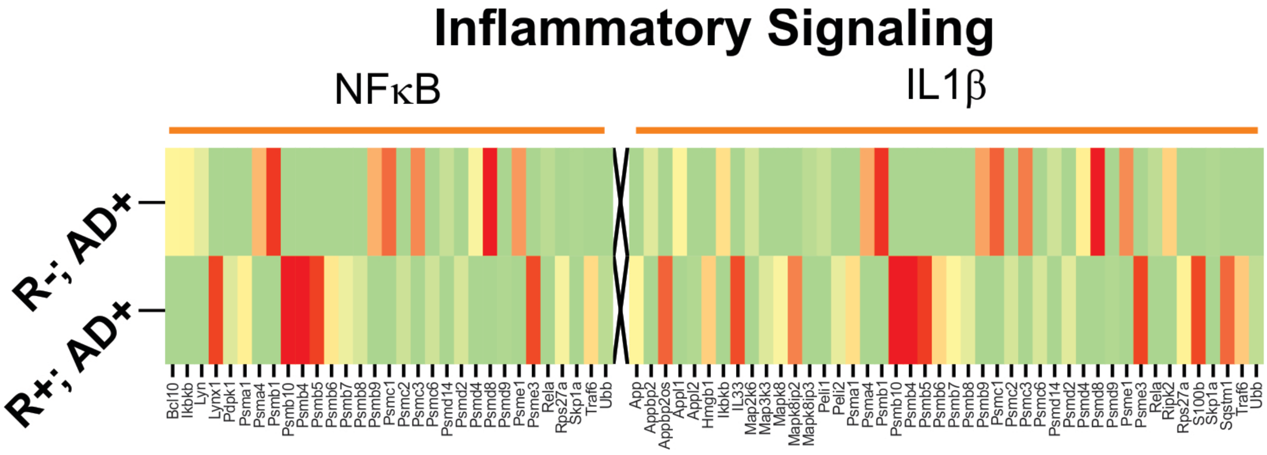

**Fig. S11: NFκB and IL1β pro-inflammatory cytokine signaling is modulated by ACE expression in microglia.** ACE modulated gene expression related to NFκB and IL1β signaling in microglia.
